# Extracellular stiffness regulates site-specific lung development

**DOI:** 10.1101/2025.01.12.632508

**Authors:** Zhiying Liao, Junjie Lv, Dong Wang, Xuepeng Chen, Jincun Zhao, Tao Xu, Hao Meng, Huisheng Liu

## Abstract

Extracellular matrix (ECM) stiffness plays a crucial role in regulating cell fate and maturation, but its influence on lung development is limited known. Here we utilized stiffness-tunable gelatin methacryloyl (GelMA) hydrogels to investigate how ECM stiffness influences site-specific lung development in a stem cell-derived lung organoid model. We found increased stiffness promoted NKX2-1+ lung progenitor cells (LPCs) generation. In airway organoids (hAWOs), stiff hydrogels directed proximal airway differentiation enriched with goblet, ciliated, and basal cells; whereas the decreased stiffness favored emergence of secretory cells in the proximal-distal transition zone and distal airway. In alveolar organoids (hALOs), increased stiffness enhanced AT2 and AT1 cells transition. Moreover, infection assays with Omicron BA.1.1 and Delta variants recapitulated the proximal-to-distal tropism of SARS-CoV-2 in the lung. Transcriptomic sequencing revealed ECM stiffness regulates lung development via Hippo, TGF-β, HIF and Wnt pathways. These findings advance mechanism understanding of ECM stiffness on lung development and provide a novel mechanical regulation for generating site-specific lung organoids.

## Introduction

The physical properties of the extracellular environment, particularly its mechanical stiffness^1^, have been demonstrated to play a crucial role in guiding tissue morphogenesis and determining cell fate^2, 3^. Extracellular matrix (ECM) stiffness varies widely across the body, ranging from the soft, pliable brain tissue with an elastic modulus of tenths of a kilopascal (kPa) to the rigid, calcified bone with a modulus reaching hundreds of megapascal (MPa)^2^. These diverse mechanical environments shape cells to adapt and function optimally within their specific tissue context. Stem cell-derived organoids serve as ideal models to study the impact of microenviroment on cell fate determination. For example, mesenchymal stem cells (MSCs) differentiate into neurogenic, myogenic, or osteogenic lineages depending on ECM stiffness^4^. Tuning mechanical stiffness of the matrix affects brain organoids growth, cellular differentiation and structural organization^5^. These findings underscore the importance of ECM mechanics in directing tissue-specific development.

The lung is a highly dynamic organ with complex structural and functional requirements that necessitate precise control of its ECM properties during development^6, 7^. Early lung development involves processes such as branching morphogenesis, vasculogenesis, and alveolar formation, all accompanied by significant changes in ECM composition and stiffness^8–10^. The lung epithelium comprises diverse cell types, with goblet and ciliated cells populating the proximal airway^11^ and secretory cells dominant in the distal regions^12, 13^. Within the alveoli, biophysical forces generated by respiratory motion maintains AT1 cell identity and loss of biophysical forces causes AT1 cells to reprogram into AT2^14^. ECM stiffness varies across different regions of the pleura where lung epithelial cells reside^15^. Advanced techniques like atomic force microscopy (AFM) have revealed regional variations in ECM stiffness, with higher stiffness in the pleura and vasculature compared to the alveolar walls^16^. Furthermore, ECM stiffness has been shown to regulate lung smooth muscle cell fate^17^. Increased ECM stiffness through photocrosslinking suppresses budding morphogenesis in embryonic lungs, highlighting the critical role of mechanical properties in lung patterning and morphogenesis^10, 18^. However, whether heterogeneous stiffness distribution suggests that localized mechanical gradients guide site-specific lung development and its underlying molecular mechanisms remain unclear.

Stem cell-derived lung organoids have been widely used to investigate lung development^11, 13, 19, 20^. These organoids are typically embedded in Matrigel for 3D structural support. However, as an ECM surrogate, Matrigel has limited mechanical tunability^21, 22^, failing to replicate the complex mechanical properties of the native lung microenvironment. In contrast, mechanically tunable hydrogels enable precise control of ECM stiffness, facilitating organoids differentiation^5, 23, 24^. Gelatin methacryloyl (GelMA), a mechanically tunable alternative to Matrigel, has been successfully used in various organoid systems, including human pluripotent stem cell (hiPSC)-derived kidney organoids^23^, intrahepatic cholangiocyte organoids^21^ and patient-derived breast cancer organoids^25^. The tunable stiffness of GelMA makes it an ideal candidate for studying ECM stiffness in directing cell differentiation during development.

This study utilized a stiffness-tunable GelMA hydrogel system to providing a 3D culture environment for human embryonic stem cells (hESCs)-derived lung organoids, aiming to investigate ECM stiffness in site-specific lung respiratory epithelial cells development. Our results revealed a correlation between ECM stiffness and lung epithelium cells fate, along with its potential underlying mechanism. This study highlights the impact of ECM stiffness on lung epithelial cells fate, offering a platform to generate region-specific lung organoids from proximal to distal to closely mimic *in vivo* lung.

## Results

### High ECM stiffness promotes lung progenitor cells (LPCs) generation through activation of Hippo and TGF-β signaling pathways

Efficient differentiation of NKX2-1+ LPCs from hESCs is critical for airway organoids (hAWOs) and alveolar organoids (hALOs) maturation^13^. Early expression of SOX2 and SOX9 play a pivotal role in airway branching morphogenesis, with SOX2 maintaining the multipotency of progenitor cells in the proximal airway, and SOX9 being more predominant in distal lung^26^. In this study, hESCs were differentiated into definitive endoderm (DE) and anterior foregut endoderm (AFE) spheroids, then encapsulated in GelMA hydrogels of varying stiffness (G30, soft; G60, intermediate; G90, stiff) to investigate stiffness-mediated effects on LPCs differentiation (Fig. 1a, Supplementary Fig. 1a and 2a).

**Fig 1.**
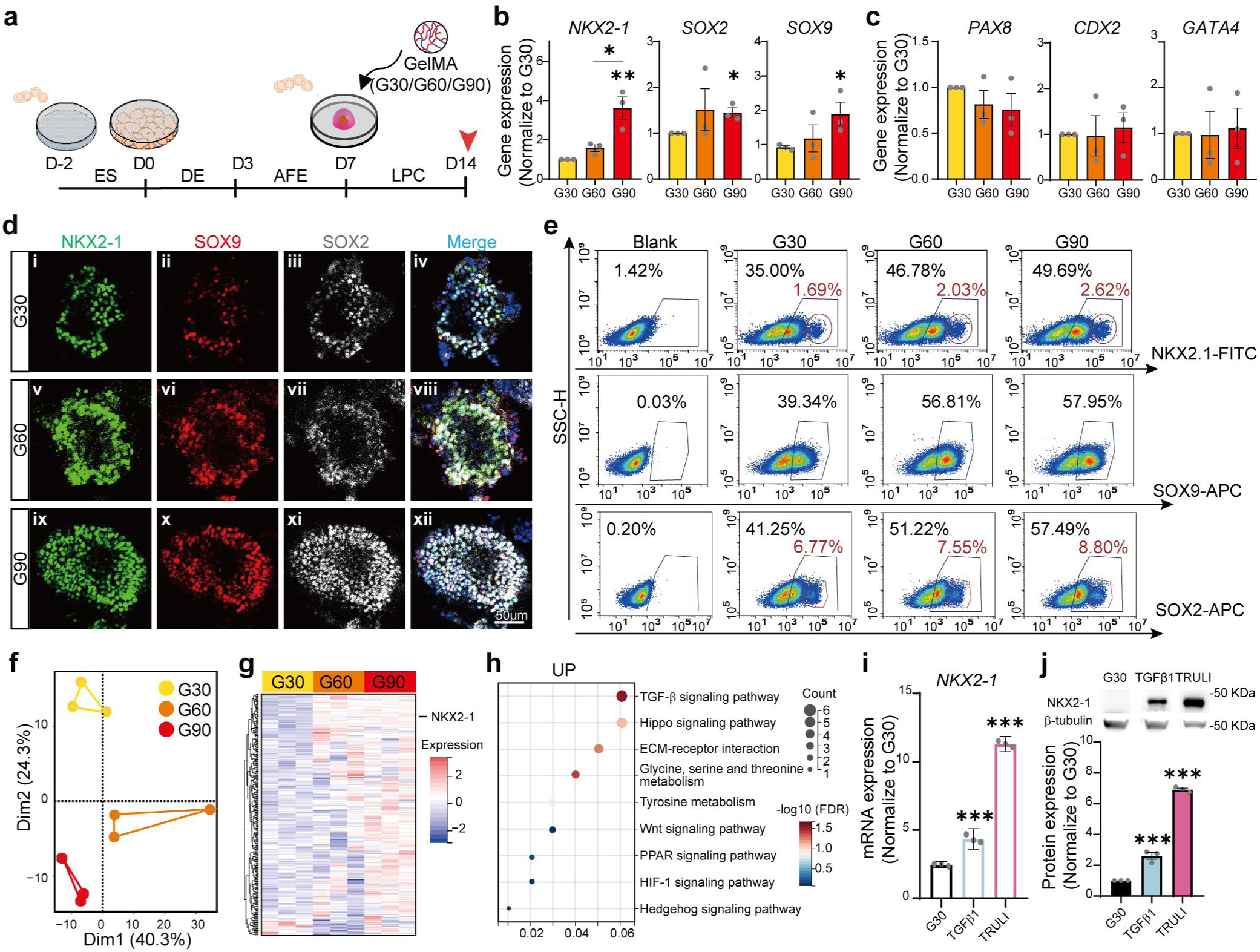
| High stiffness promotes NKX2-1+ LPCs generation by activating the TGF-β and Hippo pathways. **a**, Schematic of the directed differentiation protocol from hESCs to LPCs. Self-assembled AFE spheroids were collected at day 7 and encapsulated in 25µL GelMA hydrogels with various stiffness. ES, embryonic stem cells; DE, definitive endoderm; AFE, anterior foregut endoderm; LPCs, lung progenitor cells. Red arrow head indicates the time to collect samples. G30/G60/G90, GelMA hydrogels with low to high stiffness. **b**, Fold changes in mRNA expression of lung lineage -related genes relative to G30: *NKX2-1* (lung progenitor marker), *SOX2* (hAWOs progenitor marker); and *SOX9* (hALOs progenitor marker). **c**, Fold changes in mRNA expression of non-lung lineage genes relative to G30: *PAX8* (thyroid marker), *CDX2* (intestinal marker), *GATA4* (neuronal marker). **d**, Representative Immunofluorescence (IF) whole mounting images of lung lineage markers: NKX2-1 (green), SOX9 (red), and SOX2 (white) in LPCs differentiated in G30 (i-iv), G60 (v-viii), and G90 (ix-xii). **e**, Representative flow cytometry profiles of NKX2-1, SOX2 and SOX9 positive cells in LPCs under varying stiffness hydrogels. Blue polygon indicates positive cell population; Red circle represents the strongly positive cell population. **f**, Principal component analysis (PCA) of bulk RNA sequencing data from LPCs cultured in G30 (yellow), G60 (orange) and G90 (red). **g**, Heatmap of significantly upregulated genes, highlighting increased *NKX2-1* expression with gradient hydrogel stiffness. **h**, KEGG pathway enrichment analysis of upregulated genes. **i**, *NKX2-1* gene expression in LPCs treated with TGFβ1 (a TGF-β pathway agonist) and TRULI (a nuclear YAP activator) normalized to G30. **j**, Western blot analysis of NKX2-1 protein levels treated with TGFβ1 and TRULI. Images of IF, flow cytometry and WB are from one representative experiment from at least 3 independent experiments. For all statistical plots, data are presented as mean ± SEM, with distinct dots representing individual values from 3-5 replicates across 3 independent experiment. *p* values were calculated using ratio paired t test, one-way ANOVA and Tukey’s multiple comparison test or Dunnett’s multiple comparison test. \**p* < 0.05, \*\**p* < 0.01, \*\*\**p* < 0.001.

Hydrogel stiffness significantly enhanced *NKX2-1, SOX2* and *SOX9* expression (Fig. 1b) without promoting the expression of non-lung lineage differentiation markers like *PAX8* (thyroid), *CDX2* (intestine), *GATA4* (neuron) (Fig.1c). Immunofluorescence (IF) staining confirmed that G90 hydrogel exhibited higher NKX2-1 expression with strong co-localization of SOX2 and SOX9 (Fig. 1d). Flow cytometry further demonstrated a stiffness-dependent increase in NKX2-1+ LPCs, rising from 35.00% in G30 to 46.78% in G60 and 49.69% in G90 (Fig. 1e). Similarly, SOX9+ LPCs increased from 39.34% in G30 to 57.95% in G90 while SOX2+ LPCs rose from 41.25% to 57.49%. Notably, the G90 group exhibited a significantly higher percentage of strongly NKX2-1+ (2.62%) and SOX2+ (8.80%) cells (Fig.1e; highlighted in red circle) compared to G30 (1.69% and 6.77%, respectively) and G60 (2.03% and 7.55%, respectively).

To identify the signaling pathways mediating stiffness-regulated LPCs differentiation, bulk RNA sequencing was performed on LPCs cultured under different stiffness. The analysis revealed distinct gene expression profiles (Fig. 1f), with *NKX2-1* expression increasing consistently with stiffness (Fig. 1g). Pathways upregulated by increased stiffness were primarily associated with LPCs differentiation (TGF-β, Wnt, Hedgehog), ECM mechanics (Hippo, PPAR and Hypoxia) and metabolism (Glycine, Serine, Threonine and Tyrosine) (Fig. 1h). Importantly, apoptotic pathways were not activated, and LPCs growth, proliferation and apoptosis remained unaffected by increased stiffness (Supplementary Fig. 2b-k). Functional validation using TGFβ1 (a TGF-β pathway agonist) and TRULI (a nuclear YAP activator) in G30 hydrogel enhanced NKX2-1 expression (Fig. 1i, j), confirming the roles of TGF-β and Hippo signaling in stiffness-mediated LPCs differentiation.

### ECM mechanical gradient from stiff to soft directs proximal-to-distal hAWOs specification

hAWOs derived from LPCs typically exhibit random combinations of epithelial cells^27^, unlike the organized proximal-to-distal arrangement of airway epithelial cells observed *in vivo*^11^. The modulation of growth factors has proven insufficient to control lung cell composition, arrangement and the heterogeneity of hAWOs^28, 29^. Notably, the natural gradient in ECM stiffness, transitioning from stiff to soft across lung regions^15, 16^, inspired us to explore its role in directing lung epithelial cell fate and spatial distribution. To address this, LPCs were re-encapsulated in GelMA hydrogels with varying stiffness (G30, G60, G90) and differentiated into hAWOs (Fig. 2a). By day 49, multicellular organoids were formed and collected for further analysis (Fig. 2b).

**Fig. 2.**
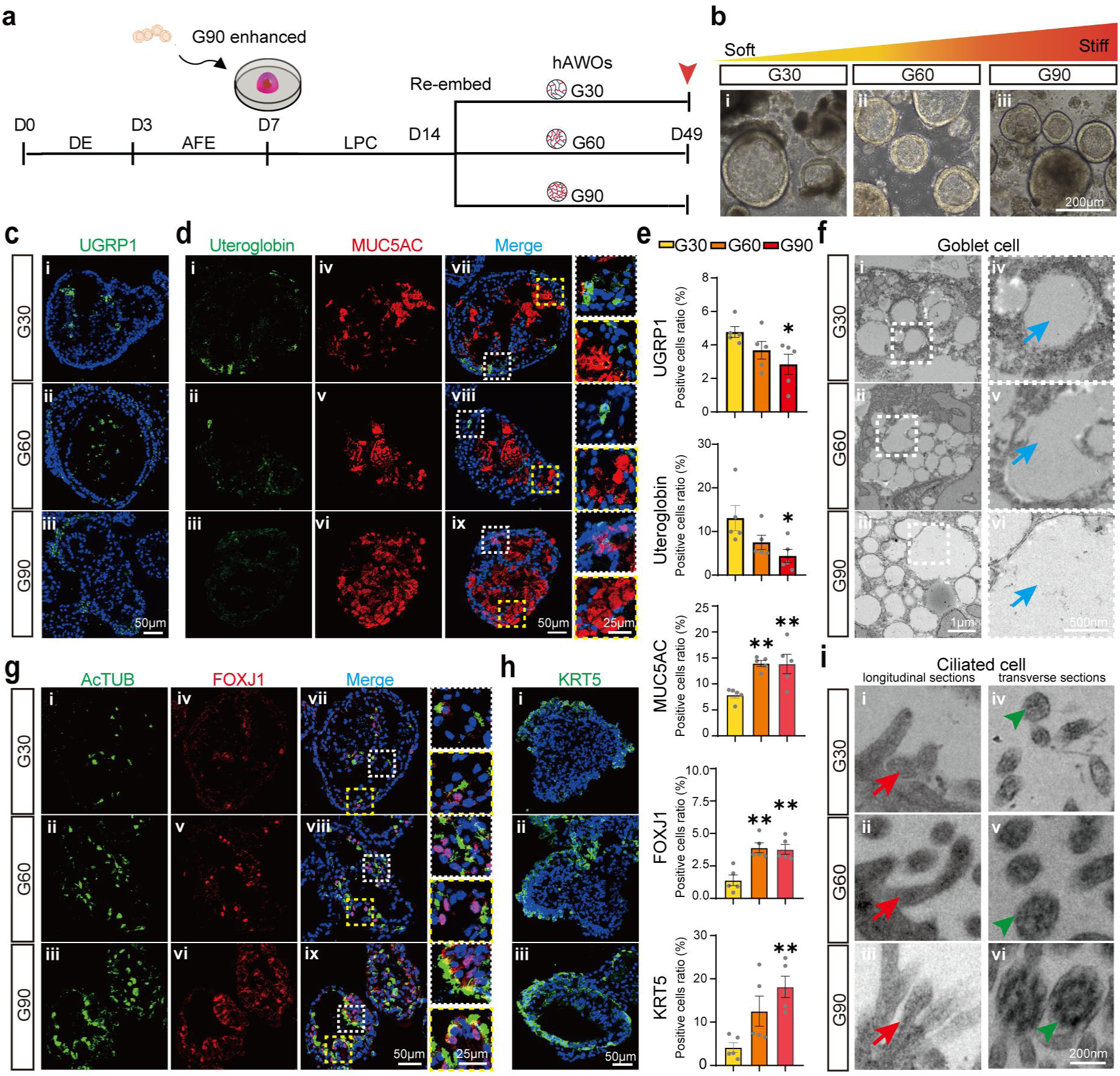
| ECM stiffness directs proximal-to-distal specification in hAWOs. **a**, Schematic of the directed differentiation protocol for hAWOs modulated by ECM stiffness. AFE spheroids were embedded in G90 at day 7 to enhance NKX2-1 expression. On day 14, LPCs were re-embedded in hydrogels of varying stiffness (G30, G60, G90) and analyzed at day 49. Red arrow indicates the time to collect the samples. **b**, Bright-field images of hAWOs cultured under different stiffness hydrogels, G30 (i), G60 (ii) and G90 (iii). **c**, Representative IF staining for UGRP1 (SCGB3A2 cell marker, green) in hAWOs cultured under G30 (i), G60 (ii) and G90 (iii). **d**, IF staining for uteroglobin (secretory cell marker, green), MUC5AC (goblet cell marker, red) in G30 (i, iv, vii), G60 (ii, v, viii), and G90 (iii, vi, ix) hAWOs. Merged images highlight regional colocalization. White dotted box highlighting uteroglobin, yellow dotted box highlighting MUC5AC. **e**, Quantification of positive cells for UGRP1, uteroglobin, MUC5AC, FOXJ1 (ciliated cell transcriptional factor), and KRT5 (basal cell marker) under varying stiffness (G30, G60, G90). High stiffness promotes goblet (MUC5AC), ciliated (FOXJ1), and basal (KRT5) cell differentiation while reducing secretory markers (UGRP1 and uteroglobin). **f**, TEM images of goblet cells under G30, G60, and G90, showing secretory vesicles (blue arrows) in organoids. **g**, IF staining for AcTUB (multiciliated cell marker, green) and FOXJ1 (red) in G30 (i, iv, vii), G60 (ii, v, viii), and G90 (iii, vi, ix), showing increased ciliated cells with stiffness. White dotted box highlighting AcTUB, yellow dotted box frame highlighting FOXJ1. **h**, IF staining for KRT5 (green) in hAWOs cultured under G30 (i), G60 (ii), and G90 (iii). **i**, TEM images of ciliated cells under G30, G60, and G90, highlighting long and dense cilia structures in longitudinal sections under higher stiffness (red arrow, left panel; i-iii), and cilia with “9+2” microtubule structures in transverse sections at high stiffness (green triangle, right panel; iv-vi). Data are presented as mean ± SEM, with distinct dots representing average values from 3–5 replicates across 3 independent experiments. *p* values were calculated using one-way ANOVA and Tukey’s multiple comparison test. \**p* < 0.05, \*\**p* < 0.01, \*\*\**p* < 0.001.

Our results revealed distinct stiffness-dependent differentiation cell fate. Secretory cells (UGRP1+ and uteroglobin+) were enriched in soft G30 hydrogel, as confirmed by IF (Fig. 2c i-iii, Fig. 2d i-iii and Fig. 2e) and flow cytometry (Supplementary Fig. 3a). Conversely, goblet cells (MUC5AC+) were more abundant in stiff G90 hydrogel (Fig. 2d iv-vi, Fig. 2e and Supplementary Fig. 3b), with more and larger mucus vesicles (blue triangle, Fig. 2f). Ciliated cells (AcTUB+ and FOXJ1+) were most prominent in G60 and G90 hydrogels (Fig. 2e, g and Supplementary Fig. 3c). Basal cells (KRT5+) were enriched in G90 from both IF and flow cytometry (Fig. 2e, h and Supplementary Fig. 3d). Notably, TEM analysis revealed that the cilia in the G90 hydrogel were longer and denser in the longitudinal section (red arrow, Fig. 2iii) and functional “9+2” microtubules were observed (green triangle, Fig. 2vi). Moreover, cilia exhibiting noticeable beating after released from G90 hydrogel (Video 1). Our results revealed by modulating ECM stiffness, region-specific hAWOs were generated: proximal-airway organoids in G90, bronchial-like organoids in G60, and distal-airway organoids in G30. These findings highlight the critical role of mechanical gradients in directing site-specific airway epithelial specification.

### High ECM stiffness promotes hALOs differentiation and maturation

The role of ECM stiffness in alveolar epithelial cells differentiation and maturation remains inconclusive. Herein, LPCs were encapsulated in GelMA hydrogels with different stiffness and differentiated into hALOs (Fig. 3a). By day 35, hALOs exhibited budding and epithelial structures (Fig. 3b).

**Fig. 3.**
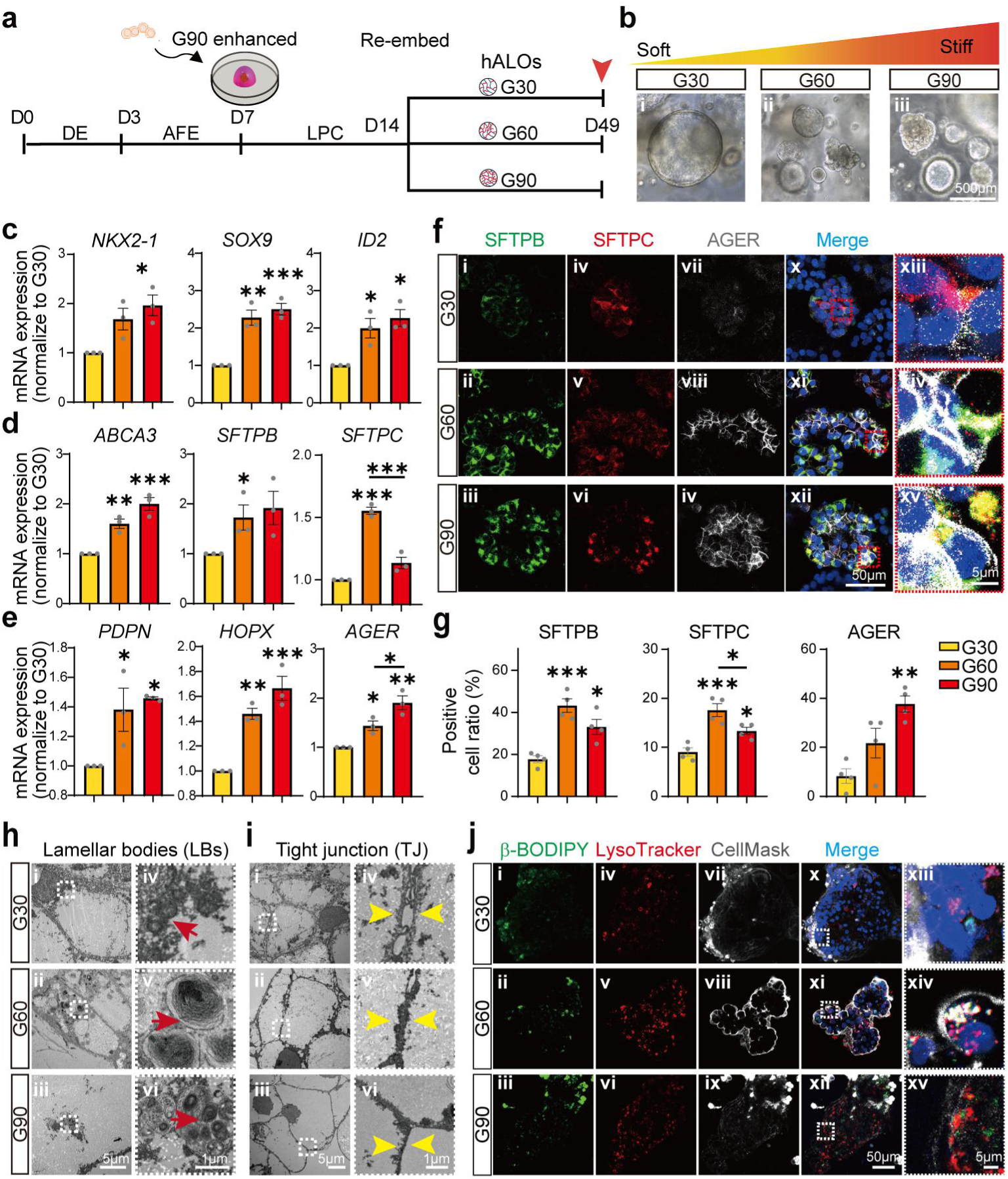
| ECM stiffness promotes alveolar epithelial cells differentiation and maturation in hALOs. **a**, Schematic illustrating stiffness-regulated differentiation of hALOs from LPCs, with sampling time points marked by red arrow head. **b**, Representative bright-field images of hALOs differentiated from G30 (i), G60 (ii) and G90 (iii). Organoids cultured in GelMA hydrogels displayed normal epithelial morphology. **c-e**, qPCR analysis of mRNA expression for (**c**) progenitor markers (*NKX2-1*, *SOX9* and *ID2*), (**d**) AT2 markers (*ABCA3*, *SFTPB*, and *SFTPC*), and (**e**) AT1 markers (*PDPN*, *HOPX*, and *AGER*). **f**, Representative IF images of SFTPB (green), SFTPC (red), and AGER (white) in hALOs. Dashed red boxes highlight magnified region from merged images. **g**, Quantification of (**f**) revealing the percentage of positive cells expressing SFTPB, SFTPC, and AGER relative to total cell nuclei. **h**, **i,** TEM images demonstrating the presence of lamellar bodies (LBs) (red arrow) in AT2 cells (**h**) and tight junctions (TJ) (yellow arrow) in AT1 cells (**i**), both are more prominent under higher stiffness. **j**, Live-cell confocal imaging of lipid metabolism and surfactant activity in hALOs cultured in G30, G60, and G90 hydrogels. β-BODIPY (green) labels neutral lipids, LysoTracker (red) indicates acidic organelles, and CellMask (white) labels the cell membrane. Merged images (x-xii) and magnified insets (xiii-xv) highlight colocalization of lipid droplets and lysosomes, indicating active lipid metabolism in AT2 cells under intermediate stiffness conditions. Stastical analyses were performed using one-way ANOVA with Tukey’s multiple comparison test; data are mean ± SEM. \**p* < 0.05, \*\**p* < 0.01, \*\*\**p* < 0.001.

The alveolar progenitor gene expressions of *NKX2-1*, *SOX9*, and *ID2* were upregulated in G90 hydrogel (Fig. 3c). The immature AT2 markers (*ABCA3, SFTPB)* were higher in both G60 and G90 while the mature AT2 marker, *SFTPC* was highest in G60 but decreased in G90 (Fig. 3d). Early AT1 markers (*PDPN*, *HOPX)* were significantly upregulated in both G60 and G90, whilst late-stage AT1 marker AGER was significantly increased in G90 (Fig. 3e). Similarly, IF staining revealed elevated SFTPB and SFTPC positive cells in both G60 (43.27% and 17.61%) and G90 (33.09% and 13.35%) compared with G30 (17.67% and 9.04%, individually), with AGER+ cells highest in G90 (37.74%) compared with G60 (21.75%) and G30 (8.27%) (Fig. 3f, g). These results indicate that higher stiffness promotes AT1 cell maturation while intermediate stiffness supports AT2 cell development.

Lamellar bodies (LBs) store and secrete pulmonary surfactant which is the key structure indicates AT2 maturation. TEM analysis showed smaller and disorganized immature LBs in G30 but larger, highly organized mature LBs in G60 and G90 (Fig. 3h). Microvilli (MV) on AT2 primarily enhance surface area and maintain alveolar homeostasis, which were observed in all three stiffness groups (Supplementary Fig. 4a). Notably, tubular myelin (TM), essential for surfactant storage and secretion in mature AT2 cells, were present in G60 and G90, which is a hallmark of mature AT2 cells (Supplementary Fig. 4b). AT1 cells in G90 displayed elongated shapes of tight junctions (TJ) compared to G30 and G60 (Fig. 3i). β-BODIPY and LysoTracker staining are powerful to study lipid droplets and lysosomal activity. We found AT2 cells under intermediate stiffness exhibited higher co-localization of β-BODIPY and LysoTracker, indicating enhanced surfactant metabolism (Fig. 3j, ii v viii). In conclusion, intermediate stiffness (G60) supports AT2 cell maturation, while higher stiffness (G90) enhances AT1 cell maturation.

### Omicron BA.1.1 and Delta variants exhibit site-specific infection in hAWOs and hALOs

Omicron BA.1.1 and Delta SARS-CoV-2 variants exhibit distinct infection tropisms in human respiratory system that Omicron preferentially infects the proximal airway while Delta has strong tropism for alveolar tissue^30^. These variants were employed to evaluate whether stiffness defined site-specific lung organoids could replicate the SARS-CoV-2 infection tropism *in vitro*. To facilitate viral infection, organoids were released from GelMA hydrogels to create the apical-out model (Fig. 4a). The epithelial layer of the organoids was altered to apical-out structure (Fig. 4b, c), with Zo-1 primarily on the exterior and E-cadherin within the organoids (Fig. 4d). hAWOs and hALOs were infected by SARS-CoV-2 variants and samples were collected at 24, 48 and 72 hours post infection (hpi) (Fig. 4e). Analysis of organoids collected at 72 hpi revealed that Omicron BA.1.1 exhibited higher infection titer in hAWOs than hALOs (Fig. 4f), whereas Delta infected more efficiently in hALOs than hAWOs (Fig. 4g). These results align with previous studies^31, 32^ showing Delta’s strong tropism for alveolar tissue and Omicron’s enhanced replication in airway epithelial cells.

**Fig. 4.**
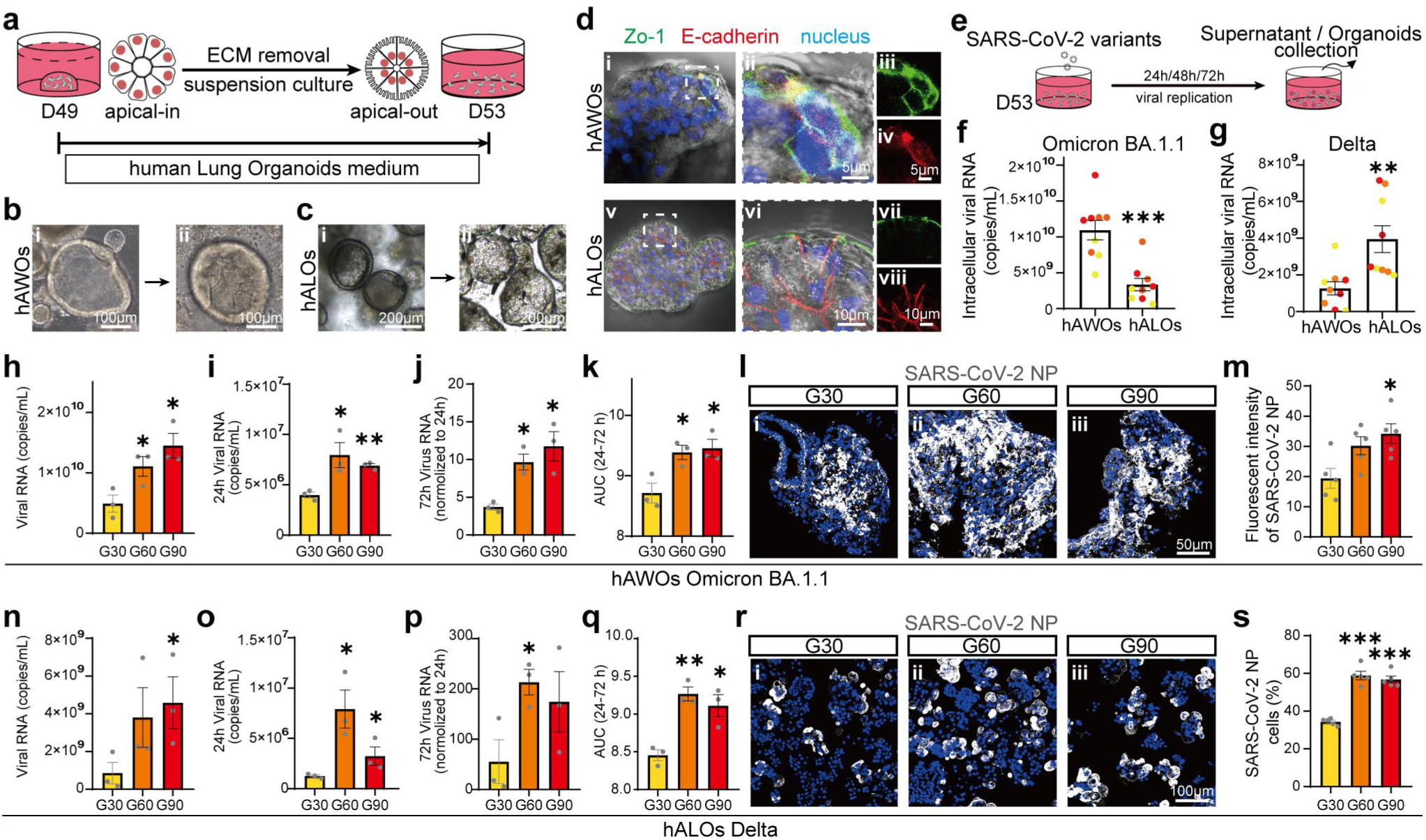
| Omicron BA.1.1 and Delta display site-specific infection tropism in hAWOs and hALOs differentiated under different stiffness. **a**, Schematic representation of construct apical-out. Apical-in organoids (D49) were transitioned to apical-out suspension culture after 3 days following ECM removal. **b**, **c**, Bright field showing the transition of apical-in organoids to apical-out orientation following ECM removal and suspension culture for hAWOs (**b**) and hALOs (**c**), individually. **d**, Representative IF staining showing TJ protein ZO-1 (green) and E-cadherin (red) in hAWOs (i-iv) and hALOs (v-viii), confirmed successful apical-out orientation. **e**, Schematic representation of the SARS-CoV-2 infection protocol in hAWOs and hALOs. On D53, organoids were exposed to SARS-CoV-2 variants (Omicron BA.1.1 and Delta) with subsequent collection of supernatant at 24, 48 and 72 hours post-infection (hpi). The organoids was lysed at 72 hpi for viral replication analysis. **f**, **g**, Intracellular viral RNA quantification in hAWOs and hALOs infected with Omicron BA.1.1 or Delta. Higher intracellular Omicron BA.1.1 loads were observed in hAWOs compared to hALOs. Whilst Delta predominantly accumulated and infected hALOs compared to hAWOs. Yellow, orange, and red dots represent G30, G60, and G90 stiffness groups, respectively. **h-k**, qPCR analysis of Omicron BA.1.1 infection in hAWOs differentiated under varying stiffness hydrogels: (**h**) intracellular viral RNA at 72 hpi, (**i**) viral RNA in supernatant at 24 hpi, (**j**) fold change of viral RNA in supernatant from 24 to 72 hpi, and (**k**) area under the curve (AUC) analysis of viral titer between 24 and 72 hpi. **l**, **m,** IF staining for SARS-CoV-2 nucleocapsid protein (NP, white) in Omicron BA.1.1-infected hAWOs cultured in G30, G60 and G90 hydrogels (**l**). (m) Quantification of (**l**) showing the percentage of NP+ cells (%) relative to the total number of nuclei. **n-p**, qPCR analysis of Delta infection in hALOs differentiated under varying stiffness hydrogels: (**n**) intracellular viral RNA at 72 hpi, (**o**) viral RNA in supernatant at 24 hpi, (**p**) fold change of viral RNA in supernatant from 24 to 72 hpi, and (**q**) AUC analysis of viral titer between 24 and 72 hpi. **r**, **s**, IF staining for SARS-CoV-2 NP (white) in Delta infected hALOs cultured in G30, G60 and G90 hydrogels (**r**). (**s**) Quantification of (**r**) showing the percentage of NP+ cells (%) relative to the total number of nuclei. For all statistical plots, data are presented as mean ± SEM, with individual points representing biological replicates (3–5 replicates, 3 independent experiments). Statistical significance were determined using unpaired t test, one-way or two-way ANOVA with Tukey’s multiple comparisons. \**p* < 0.05, \*\**p* < 0.01, \*\*\**p* < 0.001.

hAWOs differentiated in high, intermediate to low-stiffness represent proximal to distal airway. Omicron BA.1.1 infection in hAWOs differentiated under G90 had significantly higher viral loads and infection efficiency compared to soft G30 hydrogel (Fig. 4h-k), aligning with the tropism of Omicron for proximal airways. IF staining and quantification further confirmed that Omicron infection ratio were significantly higher in G90 (Fig. 4l, m). For hALOs, Delta infection efficiency was higher in organoids under high stiffness, corresponding to enhanced AT2 and AT1 cell maturation (Fig. 4n-q). IF staining and quantification further confirmed Delta infection ratio were significantly higher in the G60 and G90 groups (Fig. 4r, s). In conclusion, site-specific lung organoids derived from tunable-stiffness hydrogels effectively recapitulate SARS-CoV-2 infection tropism *in vivo*. Omicron BA.1.1 preferentially infects proximal airway organoids, while Delta targets alveolar organoids. These findings provide a valuable platform for studying SARS-CoV-2 pathogenesis and variant-specific tropisms.

### ECM stiffness determines epithelial cell fate in hAWOs via Hippo, Hypoxia, and Wnt signaling pathways

To elucidate how ECM stiffness influences epithelial differentiation in hAWOs, single-cell RNA sequencing (scRNA-seq) was performed on 31,344 cells sorted from hAWOs from varying stiffness (Fig. 5a). Cells were categorized into mesenchymal, epithelial, and neuronal lineages (Supplementary Fig. 5a, b), with epithelial cells forming the largest cluster. 31,344 cells were classified into 14 distinct subtypes (Fig. 5b, c, Supplementary Fig. 5c). Goblet cells were predominantly enriched in the G90 group, while secretory cells (Secretory1 and Secretory2) were more abundant in G30 (Fig. 5d and Supplementary Fig. 5d-i), consistent with gene- and protein-level phenotypes (Fig. 2).

**Fig. 5.**
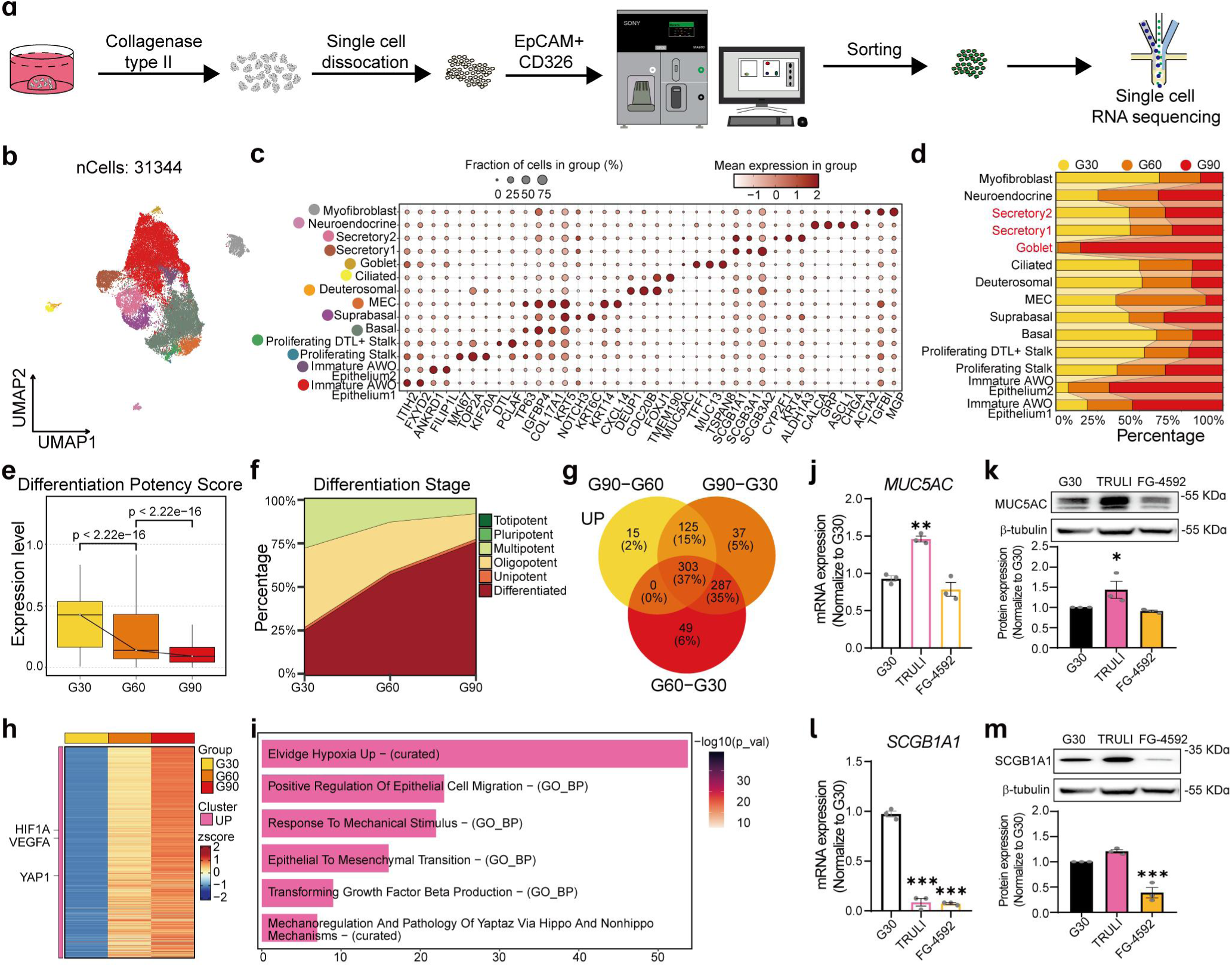
| Transcriptional results reveal ECM stiffness directs airway epithelial cell fate via regulating YAP1 and HIF1A. **a**, Workflow for single-cell RNA-seqencing (scRNA-seq) analysis of hAWOs. Cells were released from GelMA using collagenase type II, then dissociated into single cells with TrypLE, enriched via EpCAM+ sorting, and subjected to scRNA-seq. **b**, UMAP visualization of 31,344 cells were grouped into 14 distinct clusters based on curated cell type markers. **c**, Dot plot showing marker genes expressions across 14 cell clusters. The size of dots indicates the fraction of cells expressing the gene and the color of dots represents the mean gene expression, respectively. **d**, Stacked bar chart displaying the distribution of cell types across various stiffness of G30, G60, and G90. **e**, Differentiation potency scores calculated using CytoTRACE2, indicating reduced differentiation potency with increased stiffness. **f**, Predicted absolute differentiation stages based on scRNA-seq data, indicating higher stiffness supports transitions to unipotent and differentiated states (categorized as Totipotent, Pluripotent, Multipotent, Oligopotent, Unipotent, and Differentiated). **g,** Venn diagram showing upregulated genes shared across low to high stiffness. Overlapping regions highlight genes shared between each two groups (G60 vs G30, G90 vs G60 and G90 vs G30). **h**, Heatmap showing gene expression profiles of key genes (e.g., *HIF1A*, *VEGFA*, *YAP1*) regulated by higher stiffness. **i**, Pathway enrichment analysis of upregulated genes using GO-BP and curated databases, highlighting hypoxia, epithelial migration, mechanical stiffness response, epithelial-to-mesenchymal transition (EMT), TGF-β, and Hippo signaling pathways. **j**, **k**, MUC5AC expression analysis via qPCR (**j**) and western blot (**k**) in G30 organoids treated with TRULI and FG-4592 (HIF1A stabilizer). \**p* < 0.05, \*\**p* < 0.01, analyzed by one-way ANOVA with Dunnett’s multiple comparisons (n=3). **l**, **m**, SCGB1A1 expression analysis via qPCR (**l**) and western blot (**m**) in G30 organoids treated with TRULI and FG-4592. ***p < 0.001, analyzed by one-way ANOVA with Dunnett’s multiple comparisons (n=3).

Differentiation potency analysis revealed the highest scores in G90 (Fig. 5e), with approximately 70% of cells were terminally differentiated, indicating advanced maturation under high stiffness (Fig. 5f). 695 intersecting differentially expressed genes (DEGs) were identified across stiffness conditions, with 303 genes were commonly upregulated, including *HIF1A*, *VEGFA*, and *YAP1* (Fig. 5g, h). Enriched pathways included hypoxia, epithelial migration, mechanical stiffness response, epithelial-to-mesenchymal transition (EMT), TGF-β, and Hippo signaling pathways (Fig. 5i), while 353 overlapping downregulated genes were associated with cell metabolism and Wnt signaling pathways (Supplementary Fig. 5j-l).

To validate the roles of Hypoxia and Hippo pathways in hAWOs cell fate, FG-4592 (a HIF1A stabilizer) and TRULI were applied to G30 hAWOs. TRULI significantly increased MUC5AC expression by 1.5-fold while FG-4592 alone had no effect (Fig. 5j, k). Interestingly, combined TRULI and FG-4592 significantly amplified MUC5AC expression by 4-fold (*P*<0.001) (Supplementary Fig. 5m). Both TRULI and FG-4592 suppressed *SCGB1A1* and *SCGB3A2* expression (*P*<0.001) by 5- and 10-fold individually, while the combination yielding a more pronounced reduction of 50- and 25-fold respectively (Fig. 5l and Supplementary Fig. 5m, n). Western blot confirmed FG-4592 reduced SCGB1A1 protein levels (Fig. 5m). These results suggest that the synergistic action of Hippo and Hypoxia pathways more effectively directs hAWOs epithelial cells towards the proximal airway. Wnt pathway activation with CHIR-99021 suppressed *MUC5AC* (*P*<0.01) but increased *SCGB3A2* (*P*<0.001) expression in G30 (Supplementary Fig. 5o). In summary, ECM stiffness predominantly regulates epithelial cell fate in hAWOs via Hippo, Hypoxia, and Wnt signaling pathways, enabling the generation of site-specific hAWOs through stiffness modulation.

### ECM stiffness promotes hALOs maturation via Hippo and Wnt signaling pathways

I*n vitro* experiments revealed that increased stiffness regulates AT2 and AT1 cell fates with intermediate stiffness favoring AT2 and stiff hydrogel preferentially promoting AT1 (Fig. 3). To elucidate the underlying mechanisms, scRNA-seq on 4,030 hALOs cells across varying stiffness (G30, G60, and G90) were categorized into epithelial, mesenchymal, and neuronal lineages and further annotated into 20 subtypes, including undifferentiated epithelium, tip ETV5+, AT2, AT1, and proliferating AT2 and AT1 cells (Fig. 6a, b and Supplementary Fig. 6a-c). AT2 cells were most abundant in G60, whereas AT1 cells were more prevalent in G90 (Fig.6c), consistent with experimental observations (Fig. 3). Further differentiation potency analysis and pseudotime trajectory inference via Slingshot demonstrated two distinct lineages from tip ETV5+ progenitors to mature AT1 and AT2 cells (Fig. 6d, e; Supplementary Fig. 6d, e). Proliferating AT1 cells co-expressed mature AT1 (*CLIC5*, *CAV1*, *AGER*) and AT2 (*ABCA3*, *LAMP3*, *SFTPC*) markers, suggesting the transition between proliferating AT2 and proliferating AT1 (Fig. 6b). YAP is widely recognized for its critical role in the differentiation and regeneration of alveolar epithelium cells^33^, therefore we investigated its expression in different alveolar epithelial cells across stiffness groups. YAP1 expression increased with stiffness in tip ETV5+, AT2, proliferating AT2 cells and AT1 (Fig. 6f). IF staining confirmed that YAP signaling translocated from the cytoplasm to the nucleus with increasing ECM stiffness (Fig. 6g-h). Specifically, YAP was primarily localized in the cytoplasm and colocalized with SFTPC represented AT2 cells (Fig. 6g x-xii). While YAP was transitioned from cytoplasm to nuclear localization of AT1 (AGER) under higher stiffness (Fig. 6h x-xii).

**Fig. 6.**
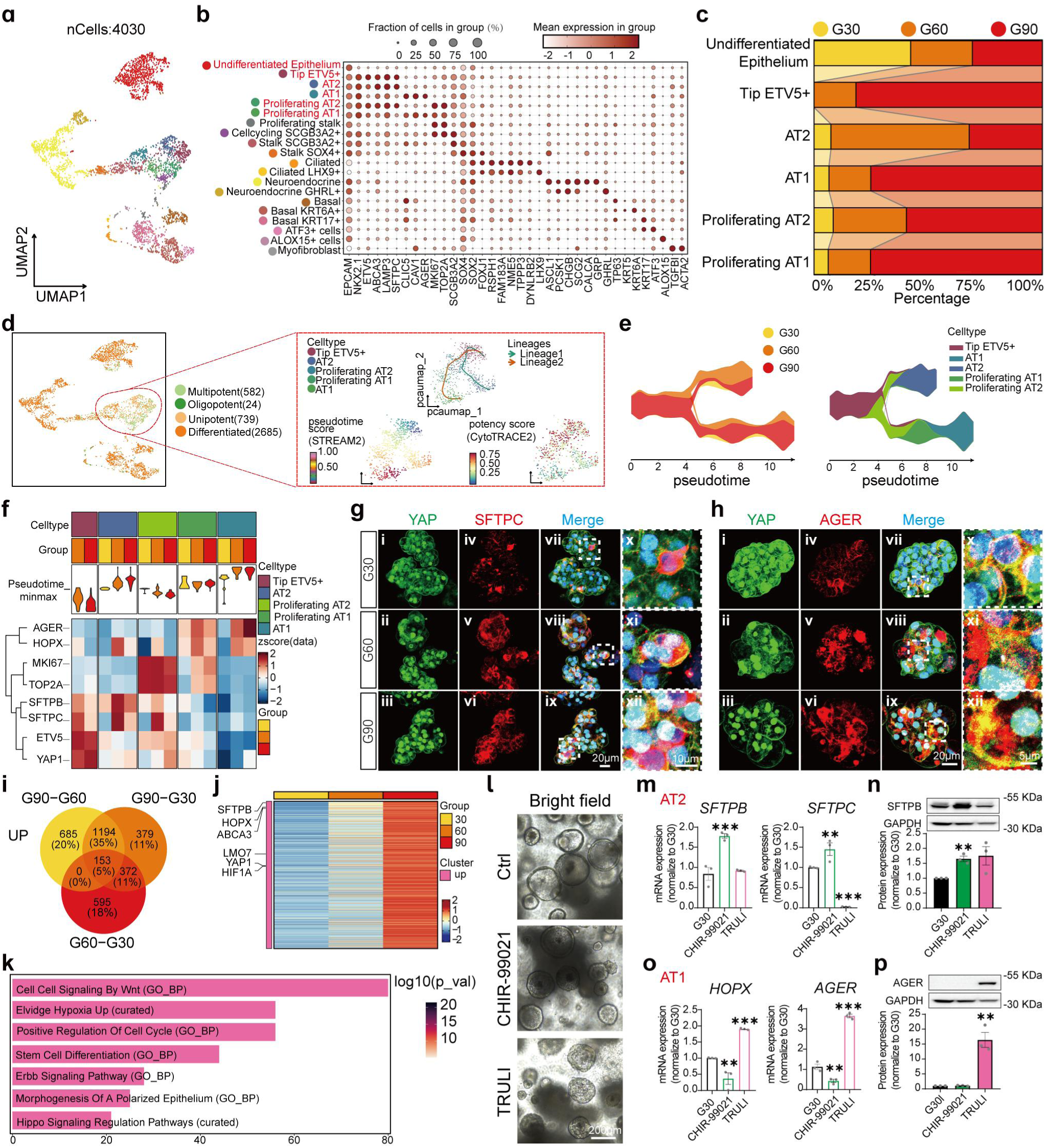
| ECM stiffness regulates hALOs differentiation and maturation via Hippo and Wnt signaling pathways. **a**, UMAP visualization of single-cells from hALOs differentiated under G30, G60 and G90, with 20 curated annotated cell types. **b**, Dot plot showing scaled marker gene expressions across 20 cell types, with dot size representing the fraction of positive expression cells and color representing mean gene expression, respectively. **c**, Stacked bar chart displaying the distribution of cell types under different stiffness. Higher stiffness promotes differentiation into maturer alveolar cells, including both AT1 and AT2. **d**, Differentiation potency analysis showing alveolar cell types are primarily multipotent. The enlarged red rectangle showing the re-computed UMAP of alveolar differentiation related cells (tip ETV5+, AT1, AT2, proliferating AT1 and proliferating AT2). Slingshot analysis visualizing 2 separate differentiation trajectories along pseudotime, with lineage bifurcations into AT1 and AT2 populations. Differentiation potency score decreases accordingly along with predicted pseudotime. **e**, STREAM trajectories of cell differentiation grouped by stiffness (left) and cell type (right), revealing stiffness-dependent shifts in differentiation dynamics, which also demonstrate there are 2 distinct branches from tip ETV5+ to 2 mature alveolar cell types. **f**, Heatmap showing dynamic expressions of key genes are unsupervised clustered into groups (e.g. *AGER* and *HOPX* for AT1, *MKI67* and *TOP2A* for proliferating cell, *SFTPB* and *SFTPC* for AT2, *ETV5* for tip ETV5+, *YAP1* for AT1 fate determination) and their correlation with pseudotime and stiffness. **g**, **h,** Representative IF staining for YAP (green) with AT2 marker SFTPC (red) (**g**) and AT1 AGER (red) (**h**) in hALOs under G30, G60, and G90. Magnified regions highlight colocalization of YAP and SFTPC or AGER. IF staining indicated YAP signaling transferred from cytoplasm to nuclear with increased ECM stiffness **i**, Venn diagram showing upregulated differential expressed genes shared with G60 vs G30, G90 vs G60 and G90 vs G30. A total of 153 genes were upregulated simultaneously with increasing stiffness. **j**, Heatmap of stiffness-upregulated genes, including *SFTPB*, *HOPX*, *ABCA3*, *LMO7*, *YAP1*, *HIF1A* across low to high stiffness. **k**, Pathway enrichment analysis of upregulated genes, highlighting alveolar differentiation related, such as Wnt, Hypoxia, and Hippo signaling pathways, along with cell cycle regulation and stem cell differentiation. **l**, Bright-field images of hALOs treated with CHIR-99021 (Wnt pathway activator) and TRULI. **m**, **n**, Expression analysis of AT2 markers (SFTPB and SFTPC) via qPCR (**m**) and western blot (**n**), showing enhanced AT2 expression with CHIR-99021 treatment. \*\**p* < 0.01, \*\*\**p* < 0.001, analyzed by one-way ANOVA with Dunnett’s multiple comparisons (n=3). **o**, **p**, Expression analysis of AT1 markers (HOPX, AGER) via qPCR (**o**) and western blot (**p**), demonstrating enhanced AT1 expression with TRULI treatment. \*\**p* < 0.01, \*\*\**p* < 0.001, analyzed by one-way ANOVA with Dunnett’s multiple comparisons (n=3).

DEGs identified 153 genes consistently upregulated and 125 downregulated across higher stiffness (Fig.6i and Supplementary Fig. 6f). Gradient-variant genes were enriched in the pathways associated with alveolar differentiation, such as Wnt, Hypoxia, and Hippo signaling pathways, along with cell cycle regulation and stem cell differentiation (Fig. 6j, k). While the down-regulated DEGs were predominantly enriched in cellular stress, oxidative phosphorylation pathways, suggesting that organoids were under stress without sufficient ECM support (Supplementary Fig. 6g, h).

Functional validation with CHIR-99021 treatment of hALOs in G30 indicated the organoids maintained the lumen structure (Fig. 6l) and significantly upregulated AT2 marker (Fig. 6m, n). While the addition of TRULI induced aggregated clumps of cells lacking visible lumen (Fig. 6l), previous study reported^34^ the correlated morphology emerged with AT1. qPCR and WB confirmed TRULI predominantly promoted AGER (4-fold at gene level and 15-fold at protein level) expression(Fig. 6o, p). In summary, ECM stiffness promoted hALOs maturation primarily through Hippo and Wnt signaling. Wnt upregulation enhances AT2 expression while inhibiting AT1 differentiation, whereas Hippo pathway promoted AT1 expression at the expense of AT2 expression.

## Discussion

In this study, a stiffness-tunable GelMA hydrogel system was employed to provide a 3D culture environment for lung organoids, enabling the investigation of ECM stiffness in site-specific lung development. Results revealed that increased stiffness upregulated NKX2.1 expression, proximal airway epithelial cells generation and function (e.g., goblet cells), alveolar maturation, and AT2/AT1 cell transition, while lower stiffness favored distal bronchial epithelial cells (e.g., secretory cells). TEM imaging showed more functional epithelial cells in higher-stiffness hydrogels, including ciliated cells with denser, longer cilia, goblet cells with larger vesicles, mature AT2 cells with well-formed LB, and AT1 with more well-developed TJ morphology. Organoids derived from stiffness-tunable hydrogels also modeled SARS-CoV-2 variants infection tropism, with Omicron BA.1.1 targeting proximal airways and Delta infecting alveolar cells. Transcriptomic data and small-molecule testing highlighted Hippo, Hypoxia, Wnt, and TGF-β as key signaling pathways. This study demonstrates how ECM stiffness determines lung epithelial cell fate, providing a platform to generate site-specific lung organoids that closely mimic the *in vivo* lung.

Embryonic stem cells differentiation into organoids at specific sites is regulated not only by small molecules but also by the composition and stiffness of the 3D culture ECM hydrogel^28^. Although Matrigel is the most widely used ECM hydrogel, its composition and mechanical properties are not adjustable^21, 22^. Alternative matrices have been explored for culturing lung organoids derived from stem cells or lung tissues. For example, polyethylene glycol (PEG) hydrogel modified with peptides has shown to support mouse lung organoids alive for 7 days^35^. However, the lack of nutrients in this synthetic hydrogel limits its suitability for long-term lung organoids culture. Recently, Zhang et al.^36^ demonstrated that incorporating hyaluronic acid (HA) into Matrigel enhances its mimicry of the natural lung. This modification accelerates lung organoid development and activates mechanotransduction pathways. While this study improved the Matrigel composition by addition of HA, the effects of more refined stiffness gradients remain unexplored. Our results clearly indicated by precisely controlling ECM stiffness, we could direct lung epithelial cell differentiation to better recapitulate *in vivo* lung epithelial organization and function.

The mechanical microenvironment regulates cell fate through interactions with downstream signaling pathways^37^. scRNA-seq of our organoids revealed ECM stiffness modulates the Hippo pathway throughout lung development involve determining lung progenitors, airway and alveolar cell fates (Supplementary Fig. 7). Hippo signaling pathway, a well-established mechanotransduction pathway that mediates its effects primarily through YAP/TAZ signaling^38, 39^. The role of Hippo signaling in human early lung development and airway cell fate is poorly understood compared to animal models. Yap-deficient mice still expressed strong NKX2.1 which indicated YAP did not directly damage lung cell identity^40^. Moreover, prior animal studies showed YAP/TAZ deletion promoted goblet cell differentiation and mucin hypersecretion^41, 42^. However, our human stem cell derived LPCs model indicated stiffness boosted NKX2-1 expression (Fig. 1) and enhanced goblet cell differentiation via activating *Yap1* (Fig 5j-m). The discrepancy between mice model and human stem cell-derived organoid model implied *in vivo* animal data may not fully reflect what occurs in human. Hippo signaling is also crucial for alveolar development, maturation and function. Inactivation of Hippo signaling or loss of YAP/TAZ impairs alveolar formation and differentiation^43, 44^. Besides, Yap mutants show increased AT1 and decreased AT2 proportions^45^. Consistently, our results confirmed that under high stiffness, YAP nuclear translocation drives AT1 differentiation in hALOs (Fig. 6f-h) and increases in AGER and HOPX expression while reducing SFTPC levels. Overall, Hippo signaling plays a critical role in determining LPCs, goblet, AT2 and AT1 cells fate, providing new insights into ECM-mediated regulation of human lung development and fate determination.

Hypoxia signaling was upregulated in high-stiffness hydrogels, likely due to the dense network hinder oxygen exchange. Previous studies demonstrated that HIF1 promotes goblet cell hyperplasia and MUC5AC expression^46, 47^. Consistently, our findings support hypoxia cooperated with Hippo signaling to promote goblet cell differentiation. Wnt signaling, GSK3β inhibitor CHIR-99021 directs distal lung patterning^48^, while preventing airway cell formation^49^. Therefore, CHIR was added to the alveolar medium to promote alveolar development. Our results revealed site-specific stiffness-dependent regulation of Wnt signaling: high stiffness downregulated Wnt signaling in hAWOs but upregulated it in hALOs. Hence, in hAWOs, our stiff hydrogel which mimic proximal ECM stiffness attuned Wnt signaling to maintain the proximal epithelial cell fate. Attenuation of the Wnt signaling rescued goblet cell differentiation was also reported in previous study^50^. In alveolar cells, Wnt signaling could promote AT2 cells generation, however, withdrawal of Wnt signaling advances the formation of mature LBs in AT2 cells^51^. We observed increased AT2 expression and LBs in intermediate stiffness hydrogels, further increased stiffness hinder AT2 development (Fig. 3d, g, h, j). In our hALOs map, we identified proliferating AT2 and proliferating AT1 cells, which ultimately adopted distinct epithelial cell fates (Fig. 6d,e and Supplementary Fig. 6d, e). This may explain the increased AT2 generation accompanied with decreased AT1.

This study investigated the generation of site-specific lung organoids through modulation of ECM stiffness and its potential mechanism. Specifically, stiff hydrogel promotes the LPCs differentiation through Hippo and TGF-β signaling (Supplementary Fig. 7a). While stiff hydrogel promotes differentiation of hAWOs enriched with goblet, ciliated, and basal cells, resembling the upper airway regions (Supplementary Fig. 7b), the soft hydrogel facilitates the differentiation and maturation of secretory cells, resembling the distal airway. Hydrogel with intermediate stiffness produces epithelial cells representing a transitional zone between proximal and distal airways. For hALOs differentiation, increased stiffness enhances alveolar maturation and modulates the AT2 and AT1 cells transition (Supplementary Fig. 7c). In conclusion, these findings offer valuable insights into human lung development and create opportunities for modeling respiratory diseases and testing therapeutic interventions.

## Materials and Methods

For details protocols and additional information, refer to the Supplementary Materials and Tables.

### Ethics statements

All experiments involving hESCs were conducted in accordance with institutional and ethical guidelines.

### GelMA hydrogel preparation and characteristics

GelMA hydrogels with varying stiffness levels were prepared by altering the methacrylation (MA) substitution degree while maintaining a constant precursor concentration (8% w/v) and crosslinking time (30s). GelMA kits with substitution degrees of 30%, 60%, and 90% were obtained from Engineering for Life Co. (EFL, Cat#EFL-GM-30, Cat#EFL-GM-60, Cat#EFL-GM-90). Hydrogel was prepared following the manufacturer’s protocol. The protocols about hydrogel preparation, reology, compression testing, swelling ratio analysis, scanning electron microscopy (SEM) and degradation testing were detailed depicted in supplementary materials and methods.

### Maintenance of hESCs

The hESCs line H1 (WiCell Research Institute) was maintained on Matrigel (Biocoat, Cat#354277) coated 6-well culture plates in mTeSR1 medium (StemCell Technologies, Cat#85850). Cells were passaged every 4 to 6 days using 0.5mM EDTA (Invitrogen, Cat#AM9261) at split ratios of 1:20 to 1:40. For differentiation experiments, hESCs were dissociated into single cells with Accutase (StemCell Technologies, Cat#07922) and replated at a density of 1.2 × 10^5^ cells/well in 24-well culture plates.

### Differentiation of hESCs into AFE spheroids

Differentiation into AFE spheroids were carried out following previously published protocols^13, 20, 52–54^ with slightly modifications. Briefly, H1 cells at ∼80% confluence were treated with DE medium for 3 days, following by incubation in AFE medium. Spheroids were formed and floated in the culture after 3 to 5 days with treatment of AFE medium.

### Encapsulation of AFE spheroids in GelMA hydrogel and differentiation into LPCs

AFE spheroids were encapsulated into GelMA hydrogel to support the 3D culture as described previously^20, 55^. GelMA were purchased from Engineering for Life Co. and the hydrogel precursor was prepared following user manual and was detailed depicted in the supplementary materials. Approximately 200 spheroids were mixed with 25 µL GelMA precursor and placed in the middle of one well of a 24-well plate. The 24 well plate was gently inverted to avoid the precipitation of spheroids and immediately crosslinked with 405 nm ultraviolet light (25mW/cm2, Curing surface light, EFL, Cat#EFL-LS1602) for 30s. Solidified GelMA domes were overlaid with LPCs basal medium to replace the photoinitiator. After 10 minutes, the medium was refreshed and replaced every other day for 7 days.

### Differentiation of LPCs into human lung organoids (hAWOs or hALOs)

LPCs were released from GelMA hydrogels using 100 μg/mL Collagenase Type II (Sigma-Aldrich, Cat#C2-28-100mg) and re-encapsulated following the same GelMA encapsulation protocol. LPCs were incubated in hAWOs or hALOs medium to obtain hAWOs and hALOs individually^56^. The compositions of hAWOs and hALOs medium were provided in supplementary materials.

### RNA extraction and qPCR analysis

The total RNA extraction of LPCs and human lung organoids, cDNA synthesis, and qPCR reactions were performed according to the manufacturer’s protocol (Qiagen, Cat#74106; Thermo Fisher Scientific, Cat#EP0753). GAPDH gene served as the housekeeping genes for normalization. Primer sequences are provided in Supplementary Table 1. Data analysis was conducted using Graphpad Prism Software 8.0.1 and normalization was performed against G30 using ΔΔCt or relative to GAPDH using ΔCt^13^.

### IF staining

IF staining was carried out following the published protocols. Briefly, hAWOs were released from GelMA and carried out following cryosection methods^57^. Whilst, the LPCs and hALOs with smaller size were released from GelMA and performed whole-mounting staining^58^. Antibodies information and dilutions were listed in Supplementary Table 2.

### Flow cytometry and fluorescence-activated cell sorting (FACS)

Flow cytometry was conducted using a Agilent Novocyte Advanteon (Agilent Technologies), and FACS was performed with a Sony MA900 sorter (Sony, Cat#LE-MA900FP). LPCs or human lung organoids were enzymatically dissociated into single cell suspensions using TrypLE (Thermo Fisher, Cat#12605028) and stained with antibodies specific for flow cytometry or FACS. Antibodies details are provided in Supplementary Table 2.

### Western blotting

Total protein was extracted from LPCs and HLOs using RIPA lysis buffer (Beyotime, Cat#P0013B) supplemented with a protease inhibitor cocktail (Beyotime, Cat#P1006). Protein concentrations were quantified using a BCA Protein Assay Kit (Beyotime, Cat#P0010). Equal amounts of total protein (30 μg) were loaded onto 4–20% precast polyacrylamide gels (GenScript, Cat#M42015C) for separation. Antibodies and their respective dilutions are listed in Supplementary Table 2. Protein molecular weight markers (GenScript, Cat#M00624-250; Invitrogen, Cat#26619) were added to determine the size of the protein bands. The intensity of Western blot bands was quantified using densitometry analysis. After acquiring the blot images, the grayscale values of the bands were measured using imageJ software.

### Transmission Electron Microscopy (TEM)

Human lung organoids were collected for TEM analysis following previously described methods^56^. Ultra-thin sections (approximately 90 nm) were cut on Ultra-microtome (Leica EM UC7), mounted on grids and stained with uranyl acetate and lead citrate. Images were capture during a digital camera integrated with a TEM (Thermo Fisher Scientific, Talos L120C G2).

### Bulk RNA sequencing

LPCs from G30, G60 and G90 groups were lysed with TRIzol^TM^ reagent (Thermo Fisher, Cat# 15596018) and flash frozen in liquid nitrogen for bulk RNA sequencing. Quality testing, database construction and RNA sequencing were subsequently performed by Annoroad Gene Technology (Beijing China) (three typical samples were taken from each group for mapping analysis). A gene expression matrix (tpm) of 60625 genes was generated. The initial exploratory analysis included PCA, heatmaps and bubble plots, were performed using the SangerBox online platform (http://www.sangerbox.com/tool)^59^. Differential expression analysis identified significantly upregulated genes with FDR<0.05 and |log_2_(FC)| >1. Bubble plots highlighted key pathways enriched in KEGG and other pathways of interest.

### scRNA-seq and analysis

Single-cell suspensions were prepared from human lung organoids differentiated under varying stiffness. Quality control, RNA sequencing, and library construction for hAWOs were performed using the C4 platform provided by Geneplus Gene Technology. The raw sequencing reads for hAWOs were aligned to the GRCh38 genome, yielding a total of 77,778 reads from 25,926 cells. Stressed cells from organoids were filtered using Gruffi (v1.5.5). Downstream analyses were performed using Seurat (v5.1.0). A total of 14 unique cell types were annotated for hAWOs. For hALOs, these processes were carried out using the 10x Genomics platform provided by Annoroad Gene Technology.Similarly, raw reads for hALOs were mapped to the GRCh38 genome as well, resulting in 407,789 reads from 8,132 cells, and 20 unique cell types were identified for hALOs.

Initial exploratory analyses included Venn diagrams, heatmaps, and pathway enrichment visualizations based on GO-BP and curated databases. CytoTRACE2 (v1.0.0) was employed to predict cell potency categories and absolute developmental potential from organoids scRNA-seq data. For alveolar trajectory inference, Slingshot (v2.12.0), STREAM (v1.1) and Partition-based Graph Abstraction (PAGA) were simultaneously applied to verify consistent trajectories based on gene expression matrix. All statistical analyses for violin and box plots were conducted using the “wilcox.test (alternative = ‘two-sided’)” function in the R package “stats.”

### Validation of regulatory pathways found in transcriptomics using small molecules

LPCs: To evaluate the role of the TGF-β and Hippo signaling pathways in NKX2-1+ LPC generation, LPCs were treated on the first day of the LPCs differentiation stage with 10 µg/mL TGFβ1 (Procell, Cat#PCK091) or 10 µM TRULI (MCE, Cat#HY-138489), while LPCs cultured in standard LPC medium served as the control. Samples were collected after 7 days for further analysis.

Human lung organoids: To examine the regulatory effects of hypoxia, Hippo, and Wnt signaling pathways on the directional differentiation and maturation of human lung organoids, day 49 organoids were treated with one of the following: 50 µM FG-4592 (MCE, Cat#HY-13426); 3 µM Chir-99021 (MCE, Cat#HY-10182); 10 µM TRULI; or combinations of these conditions. G30 lung organoids cultured in standard differentiation medium served as the control group. Treated organoids were collected after 7 to 14 days for subsequent analysis.

### SARS-CoV-2 infection

SARS-CoV-2 (Omicron BA.1.1 and Delta variants) was isolated from COVID-19 patients in Guangdong, China, by the Guangdong Provincial Center for Disease Control and Prevention. All SARS-CoV-2-related experiments were conducted in Biosafety Level 3 (BSL-3) laboratories at the Guangzhou Customs District Technology Center (Guangzhou International Bio Island, Guangzhou, China). The polarity of organoids was reversed to an apical-out orientation by culturing them in suspension before SARS-CoV-2 infection as described previously^32^. SARS-CoV-2 infection organoids was performed in human lung organoids medium at an multiplicity of infection (MOI) = 1 for 2 hours at 37 ℃. Then, the organoids were washed three times with DPBS. Fresh HLO culture medium was added and incubated at 37 ℃ for 72 h. Supernatant was collected at specified time points (24, 48, 72 hpi) for viral titer determination. After 72 hours, the organoids were collected for qPCR and IF staining.

### Viral RNA isolation and quantification of viral titers

Viral RNA was isolated from both the organoids and supernatant of infected organoids following previously published methods with slight modifications^60^. The pcDNA3.1 plasmid containing SARS-CoV-2 N gene fragments (4049 bp) was synthesized in Dr. Jincun Zhao’s Laboratory (Guangzhou National Laboratory, Guangdong, China) and served as a positive control for viral quantification. The complete map of the insert is provided in Supplementary Data 1. The plasmid was constructed using EcoRI and XhoI restriction sites, and the sequence was validated by Sanger sequencing. Purified plasmid DNA was quantified, and DNA copy numbers were calculated based on the virus genome equivalents formula^61^. A stock solution of 10^10^ copies/uL was prepared, and 10-fold serial dilutions were performed to obtain 10^-2^ to 10^-6^ copies/μL working solutions. Standard qPCR reactions were performed using primers specific to SARS-CoV-2 (listed in Supplementary Table 1). ΔCt values obtained from the qPCR of serial dilutions were plotted to construct the standard curve. The viral titer in each sample was then calculated by inputting its ΔCt value into the equation derived from the standard curve. AUC was calculated from the viral titer from different time points indicated in the y axis.

### Image analysis

Inverted fluorescence microscope (Nikon, ECLIPSE Ts2-FL) was used to capture the bright field of the organoids. The radium of the organoids was measured by imageJ. Confocal imaging was conducted with a Carl Zeiss LSM 980 inverted confocal laser scanning microscope and Zen 2.6 software was used to generate 3D Z-stack images. Image processing and analysis were performed using imageJ software. The percentage of positive cells was calculated as the number of positive cells divided by the number of nuclei. Image analysis and quantification of intracellular mucin content in the airway, based on published literature^62^. The area and length of the stain were applied to the volume density to translate into a 3D estimate of mucin volume per square millimeter of airway epithelial.

### Experimental replicates and statistical analysis

All the experiments were repeated at least three times. The data are presented as the means ± Standard Error of Mean (SEM.). All comparisons were statistically analyzed by one-way ANOVA and Tukey’s or Dunnett’s multiple comparison test. All of the statistical analyses in this study were performed with GraphPad Prism 8.0.1 software. A *p* value less than 0.05 is considered to indicate statistical significance.

## Acknowledgements

This study was supported by grants from the National Key Research and Development Program of China (2020YFA0908200 and 2021YFA1101300), Guangzhou Laboratory Key Research Foundation (TL22-21).

## Author contributions

Z.L., H.M. and H.L. conceptualized and designed the experiments. J.L. conducted the analysis and interpretation of single-cell RNA sequencing data, as well as the preparation and revision of the related manuscript. Z.L. and H.M. carried out cell culture and molecular biology experiments, interpreted the data, and contributed to the drafting and polishing the manuscript. D.W. handled SARS-CoV-2-related experiments in the P3 laboratory. X.C., J.Z., T.X., H.M. and H.L. supervised the study, provided data interpretation and collaborated with all authors for input. All authors reviewed and approved the final version of the manuscript.

## Competing interests

The authors declare no competing interests.

## Notes

### Competing Interest Statement

The authors have declared no competing interest.

