## Supplementary Materials for "Extracellular stiffness regulates site-specific lung development"

**This PDF file includes:**

Supplementary Materials and Methods
Supplementary References
Supplementary Figure Legends
Supplementary Figures 1 to 7
Supplementary Tables 1 to 2

### Supplementary Materials and Methods

#### Preparation and characterization of GelMA hydrogel with various stiffness

##### Preparation of GelMA hydrogel with various stiffness

GelMA hydrogels with varying stiffness levels were prepared by altering the methacrylation (MA) substitution degree while maintaining a constant precursor concentration (8% w/v) and crosslinking time (30s). GelMA kits with substitution degrees of 30%, 60%, and 90% were obtained from Engineering for Life Co. (EFL, Cat#EFL-GM-30, Cat#EFL-GM-60, Cat#EFL-GM-90). Hydrogel was prepared following the manufacturer's protocol. Briefly, 8% w/v GelMA precursor was prepared by adding 0.25% w/v lithium phenyl-2,4,6-trimethylbenzoylphosphinate (LAP) photoinitiator to GelMA powder. The GelMA precursor was filtered through 0.22 µm filter (33mm, Merck Millipore, Cat#SLGP033RB-250EA) for sterilization. To prevent solidification, the hydrogel precursor was kept at 37°C on a heat plate and shielded from light until ready for cell mixing.

##### Rheological Analysis

The storage modulus (G′) and loss modulus (G″) of hydrogels were measured by small-strain oscillatory shear measurements on rheometer (Anton Paar, Car#MCR 302e). Briefly, 5 mm thick GelMA hydrogels cylinders were prepared and balanced in deionized water. The mechanical response of the hydrogel sandwiched between the parallel plates of the rheometer was recorded by performing frequency sweep (0.1–50 Hz) measurements in a 1% constant strain mode at room temperature with an angular rate of 5 rad/s.

##### Determination of compression modulus

To evaluate the impact of MA grafting ratios on hydrogel stiffness, uniaxial compression testing was conducted. Cross-linked hydrogels of varying stiffness were prepared under UV light and tested using a universal test system (EFL, Cat#T560) equipped with a 5 N load cell. Each construct (n = 3 per group) was compressed at a strain rate of 1 mm/min with a preload of 0.01 N. The compression modulus was calculated from the slope of the stress-strain curve in the range of 0.05–0.1 mm strain.

##### Swelling ratio analysis

GelMA hydrogels were incubated in 1 mL of deionized water at 37°C for 120 minutes to reach equilibrium swelling. The swollen hydrogels were weighed, and the swelling ratio was calculated using the following formula:

$$\text{Swelling Ratio} = \frac{w_t - w_0}{w_0} \times 100\%$$

Where $W_t$ is the weight of the hydrogel after 120 minutes and $W_0$ is the dry weight of the hydrogel measured after lyophilization.

##### Scanning electron microscopy (SEM) and pore size analysis

To assess pore size, GelMA hydrogels of varying stiffness were frozen in liquid nitrogen, lyophilized (HUACHEN LAB, Cat#LGJ-12) overnight. The dried hydrogel was cutted to expose the cross-section and sputter-coated with gold (Quorum, Cat#Q150r S Plus). SEM imaging was captured using a scanning electron microscope (Phenom, Cat#Pharos G2). Images were analyzed using imageJ software, and pore sizes were calculated by measuring and averaging the areas of at least 15 pores per sample.

##### Degradation testing

GelMA hydrogels were crosslinked in 24-well plates, and the initial hydrogel diameter was measured. Samples were incubated in 500 µL of 20 µg/mL collagenase type II at 37°C. Hydrogels degradation was assessed by measuring the diameter of the hydrogel for up to 1 hour. Images of the samples were captured and diameter was measured for degradation analysis.

**Differentiation of hESCs into lung organoids**

hESCs at approximately 85% confluence were dissociated into single-cell suspensions using Accutase (Stem Cell, Cat#7920) and reseeded at a density of $1.6 \times 10^5$ cells/mL onto 24-well plates pre-coated with Matrigel (Corning, Cat#354277). 10 μM Y-27632 (MCE, Cat#HY-10071) was added to mTeSR1 medium on the first day. Once the cells reached 70–80% confluence, differentiation into lung organoids was initiated by sequential administration of specific growth factors as described below.

**Definitive endoderm (DE) medium**[1-3]: DE basal medium was prepared by composition of MCDB131 (Thermo Fisher, Cat#10372019） with 1.8 mg/mL glucose (Sigma, Cat#G7528-1kg), 1.5 mg/mL sodium bicarbonate (Sigma, Cat#6014-500g), 0.4% bovine serum albumin (BSA, Proliant, Cat#69700), 1% Penicillin/Streptomycin (Pen/Strep) (Thermo Fisher, Cat#15140122), 1% GlutaMAX (Thermo Fisher, Cat#35050061) and was stored at 4°C before use.100 ng/ml Activin A (StemCell, Cat#78001) and CHIR-99021 (MCE, Cat#HY-10182) with decreasing concentration of 3, 0.1 and 0 μM for 3 days were added to DE basal medium on the day of use.

**Anterior foregut (AFE) medium**[4, 5]: The AFE basal medium was prepared by addition of Advanced DMEM/F12 medium (Thermo Fisher, Cat#12634010) with 2% B27 supplement (Thermo Fisher, Cat#17504044), 1% N2 (Thermo Fisher, Cat#17502048), 1% GlutaMAX, 1% Pen/Strep, 1% HEPES (Thermo Fisher, Cat#15630080) and stored at 4°C before use. 1 μM SAG (MCE, Cat#HY-12848), 10 μM SB-431542 (MCE, Cat#HY-10431), 200 ng/mL Noggin (MCE, Cat#HY-P7051A), 500 ng/mL FGF4 (MCE, Cat#HY-P7014) and 2 μM CHIR-99021 were added to the AFE basal medium and the medium was refreshed daily for 3 to 5 days, depending on the formation of self-assemble spheroids.

**Lung progenitor cell (LPC) medium**[5]: LPC basal medium is the same with AFE basal medium. 10 μM DAPT (MCE, Cat#HY-13027), 20 ng/mL BMP4 (MCE, Cat#HY-P7007), 10 ng/mL FGF7 (MCE, Cat#HY-P70597), 10 ng/mL FGF10 (MCE, Cat#HY-P70695), 0.5 μM retinoic acid (Sigma, Cat#R2625), 3 μM CHIR-99021 were added to LPC basal medium and LPC medium was refreshed every other day for 7 days.

**Human airway organoids (hAWOs) medium**[6]: Human lung organoids basal medium was prepared by supplementation of DMEM/F12 medium (Thermo Fisher, Cat#10565018) with 1% B27 supplement, 0.25% BSA, 0.1% ITS-X (Thermo Fisher, Cat#51500056), and 1% Pen/Strep. 50 nM dexamethasone (MCE, Cat#HY-14648), 100 nM 8-Br-cAMP (MCS, Cat#HY-12318), 100 nM 3-isobutyl-1-methylxanthine (MCE, Cat#HY-12318), 10 ng/mL FGF7 were added before using. The hAWOs medium was refreshed on Monday morning, Wednesday afternoon and Friday afternoon every week for 35 to 45 days.

**Human alveolar organoids (hALOs) medium**: To obtain hALOs medium, hAWOs medium was supplemented with 3 μM CHIR-99021 and 10 μM SB-431542. The hALOs medium was refreshed Monday morning, Wednesday afternoon and Friday afternoon every week for 35–45 days.

**Live/dead staining assay**

The viability of LPCs encapsulated in hydrogels was evaluated using a Calcein-AM/propidium iodide (PI) Live/Dead Cell Viability Assay Kit (Solarbio, Cat#CA1630) according to the manufacturer's protocol. Live cells were stained with Calcein-AM, while dead cells were stained with PI. Cell viability was quantified as the percentage of dead cells (PI-positive) relative to the total cell population.

**Immunofluorescence (IF) staining**

After releasing the organoids from the hydrogel, hAWOs were rinsed with DPBS (Gibco, Cat#14190250) twice and subsequently fixed in 4% w/v paraformaldehyde (PFA) (Sangon Biotech, Cat#E672002-0500) for 30 minutes at 18-25 ℃ . Following fixation, organoids were washed three times with DPBS and dehydrated in 30% sucrose at 4 ℃ until they settled at the bottom of the centrifuge tube. The organoids were then embedded in Cryo Embedding Medium (Biosharp, Cat#BL557A), flash-frozen in liquid nitrogen, and stored at −80 ℃. Frozen organoids were sectioned into 10 μm slices using a cryomicrotome (Minux® FS800, RWD Life science Co.,LTD) and mounted on adhesive slides for long-term storage at −20 ℃. Before staining, sections were briefly fixed again in 4% PFA, permeabilized, and blocked for 30–60 minutes in blocking buffer (Beyotime, Cat#P0260). Primary antibodies diluted in antibody diluent (Beyotime, Cat#P0103) were applied and incubated overnight at 4 ℃. Secondary antibodies and Hoechst 33342 (Thermo Scientific, Cat#875756-97-1) were added for 1h at room temperature. Finally, stained sample were mounted using anti-fading mounting medium (ACMEC, Cat#AS2100). LPCs and hALOs with small size used whole mounting without cryosection. Briefly,the samples were collected and precipitated at the bottom of a 15 mL centrifuge tube, then fixed in 4% PFA and stained following the staining protocol described above.

**Flow cytometry and fluorescence-activated cell sorting (FACS)**

Organoids were dissociated into single cells using TrypLE (Thermo Fisher, Cat#12605028) at 37°C for 5-15 minutes, with gentle pipetting every 6 minutes to minimize cell clumping. An equal volume of DMEM/F-12 medium supplemented with 10 µM Y-27632 was added to terminate the digestion. Single cells were centrifuged at 300 × *g* for 3 minutes.

For flow cytometry analysis, single cells were fixed in 4% PFA for 30 minutes. Cells were permeabilized, blocked, and stained with primary and secondary antibodies according to the IF staining protocol. Stained cells were resuspended in sorting buffer (1 mM EDTA + 1% BSA in PBS) and passed through a 70 µm cell strainer (BD Falcon, Cat#352235) into FACS tubes and protecting from light. The stained single cells were ran and analyzed using Agilent Novocyte Advanteon (Agilent Technologies).

For FACS, the dissociation protocol of organoids was the same as above except all steps were performed on ice to preserve cell viability. After digestion, cells were incubated with EPCAM antibodies (Biolegend, Cat#369810) on ice for 30 minutes. Single cells with EPCAM positive were sorted using Sony MA900 sorter (Cat#LE-MA900FP) and collected in sorting buffer with Y-27632. Data were processed with NovoExpress (Agilent Novocyte Advanteon) and FlowJo_V10 software.

**Transmission Electron Microscopy (TEM)**

Organoids were collected and fixed in 2.5% glutaraldehyde for 1 hour at room temperature, then transferred into fresh 2.5% glutaraldehyde for preservation at 4℃. Fixed samples were washed in 0.1 M PBS (PH=7.4) for 15 minutes per wash, repeating this process four times. The samples were transferred to 1% osmium tetroxide for 1.5 hours at room temperature (performed in a fume hood due to osmium toxicity). After washing with PBS for 3 times, samples were sequentially dehydrated in graded ethanol solutions (50%, 70%, 80%, 90%, and 100% × 2) for 5 minutes each, followed by dehydration in acetone.

Samples were infiltrated with a 1:1 mixture of acetone and 812 resin for 2–4 hours at room temperature, and then embedded in fresh 812 resin, polymerization at 40°C for 24 hours, and subsequently at 60°C for 12 hours. Ultrathin sections (80–100 nm) were sectioned using an ultramicrotome, mounted on grids, and stained with uranyl acetate and lead citrate, and imaged using a

TEM (Thermo Fisher Scientific, Talos L120C G2).

**Viral RNA isolation and quantification**

To isolate viral RNA from infected lung organoids, SARS-CoV-2 infected organoids were lysed in 600 µL of RLT Plus buffer (QIAGEN, Cat#1053393) supplemented with 1% β-mercaptoethanol (Gibco, Cat#21985023). The lysate was mixed thoroughly with a pipette until complete lysis was achieved. The lysate was then stored at −20°C or −80°C for subsequent RNA isolation using the QIAGEN RNeasy Plus Mini Kit, following the manufacturer's instructions (QIAGEN, Cat#74106).

Viral RNA from the supernatant was isolated by combining 140 µL of supernatant with 560 µL of AVL buffer (QIAGEN, Cat#19089). The mixture was thoroughly vortexed and stored at −20°C until processing with the QIAGEN Viral RNA Mini Kit (QIAGEN, Cat#52906) according to the manufacturer's protocol. cDNA synthesis was performed with equal amounts of RNA from each sample using the Maxima H Minus First Strand cDNA Synthesis Kit (Thermo Fisher, Cat#EP0753). DNA copy numbers were calculated using the following formula for genome equivalents:

$$\text{DNA Copies} = \frac{\text{DNA amount (g)} \times 6.022 \times 10^{23}\ \text{(copies/mol)}}{\text{Length (bp)} \times 660\ \text{(g/mol/dp)}}$$

where DNA amount is 68.8 ng/µL, $6.022 \times 10^{23}$ is Avogadro's number, length vector + insert is the transcript length in nucleotides (4049 bp), and 660 is the average molecular weight of a single nucleotide (estimated at 660 Da). This calculation enabled the determination of DNA copy numbers for subsequent analysis.

**Single-cell RNA sequencing (scRNA-seq) of organoids and analysis**

**Preparation of Single-cell Suspensions**

Lung organoids cultured under different stiffness conditions for 49 days were dissociated into single-cell suspensions using TrypLE (Thermo Fisher, Cat#12605028). The suspensions were passed through a 70 µm cell strainer (BD Falcon, Cat#352235) into FACS tubes, ensuring a single-cell yield of over 90%. For hAWOs, epithelial cell-positive populations were pre-sorted using a flow cytometer, following previously described methods. Cell viability and concentration were assessed using Trypan Blue Stain (Invitrogen, Cat#15250061) with the Countess™ 3 Automated Cell Counter (Life Technologies). Final cell concentrations were adjusted to 1000 cells/µL with a viability of ≥80%.

**Library Construction and Sequencing for hAWOs**

hAWOs were sequenced on the C4 platform by Geneplus Gene Technology. Single-cell suspensions were processed using the DNBelab™ C Series High-throughput scRNA Library Kit (MGI, Cat#940-001818-00). Libraries were constructed following the manufacturer's instructions, including cDNA synthesis, size selection, end repair, A-tailing, adapter ligation, index library amplification (PCR), and library circularization of the sequencing library to build a library of 3' transcripts. Purification and quantification was performed using the Qubit ssDNA Assay Kit (Thermo Fisher) and Qsep100 (Bioptic). Sequencing was conducted on the DNBelab™ C4/TaiM4 platform with paired-end reads on a DNBSEQ-T7 sequencer.

**Library Construction and Sequencing for hALOs**

hALOs were sequenced on the 10x Genomics platform by Annoroad Gene Technology. Single-cell suspensions were combined with Gel Beads and oil droplets in separate channels of Chromium Chip G (10x Genomics, Cat#1000121) to generate Gel Bead-In-Emulsions (GEMs) via microfluidics. Following the manufacturer's protocol, mRNA was reverse-transcribed into complementary DNA (cDNA) and subsequently amplified into double-stranded cDNA. A-tailing was performed, followed by ligation of sequencing adapters, including the Read2 primer. The resulting libraries incorporated P5 and P7 adapters and dual indices, enabling compatibility with Illumina sequencing platforms. The quality-controlled libraries were sequenced on an Illumina platform by Annoroad Gene Technology, ensuring high-quality data acquisition for downstream analysis.

**scRNA-seq data pre-processing and quality control**

Raw sequencing data was processed with the DNBelab™ C Series Single-cell RNA Analysis Report V2.1.1 (hAWOs) and CellRanger v8.0.0 (hALOs) using default parameters. Reads were mapped to the GRCh38 genome for hAWOs (25,926 cells; 77,778 reads) and GRCh38 genome for hALOs (8,132 cells; 407,789 reads). Low-quality cells were excluded based on a cutoff of fewer than 500 genes or 1000 UMIs. Doublets were predicted using DoubletFinder (v2.0.4)[7], DoubletDetection (v4.2)[8], scDblFinder (v1.18.0)[9], and Scrublet (v0.2.3)[10], and cells flagged by at least two methods were excluded. Gruffi (v1.5.5)[11] was used to remove stressed cells of organoids with default parameters. All statistical analysis for violin charts and box charts is performed using 'wilcox.test (alternative=" two-sided ")' in the R package "stats".

**Normalizing, integrating, clustering and annotation**

Downstream analysis was performed using Seurat (v5.1.0)[12]. After quality control, gene-cell count matrices for hAWOs (G30, G60, G90) and hALOs (G30, G60, G90) were merged separately using the 'Seurat::merge()' function. Data normalization and batch correction were conducted with 'SCTransform()'. Dimensionality reduction was achieved by selecting PCs with a cumulative variance higher than 80% using 'RunPCA()', followed by constructing a KNN graph with 'FindNeighbors()'. Unsupervised clustering was performed using the Louvain algorithm of 'FindClusters()' with resolutions from 0.1 to 2.0, and low dimensionality visualization was conducted via 'RunUMAP()'. Considering noise-related clusters, we further performed subclustering on each cluster with resolution from 1.0 to 2.0, clusters with valuable biological features were remained while artificial effects clusters (low-sequencing-depth, abnormal mitochondrial percent etc.) were excluded in the later analysis.

Manual annotation was performed using curated cell type markers[13-15]. Clusters were categorized into three main groups: Epithelium, Mesenchyme and Neuron. For both hAWOs and hALOs scRNA data, epithelial cells constituted the largest proportion, displaying substantial heterogeneity within their cluster. Then we re-annotated each main group through unsupervised high-resolution clustering and cell type enrichment analysis via 'AddModuleScore()'. A total of 14 cell types were identified for hAWOs and 20 for hALOs. MetaNeighbor[16] correlation analysis using 'MetaNeighborUS()' confirmed cell type replicability across stiffness conditions.

**Stiffness-mediated differentially expressed genes (DEG) analysis**

DEG analysis was performed using the 'FindAllMarkers()' function in Seurat, calculating fold change and performing Wilcoxon Rank Sum tests. DEG sets for comparisons (G30 vs. G60, G60 vs. G90, G30 vs. G90) were intersected to identify genes consistently upregulated or downregulated with increasing stiffness.

**Absolute differentiation potential score for organoid cells**

CytoTRACE2 (v1.0.0)[17] was employed to predict cellular potency categories and absolute developmental potential based on normalized gene counts. Default parameters were applied using 'cytotrace2(full_model=T)' to calculate absolute differentiation potential scores.

**Alveolar trajectory inference analysis**

To reconstruct alveolar developmental trajectories, cells from tip ETV5$^{+}$, AT2, proliferating AT2, AT1, and proliferating AT1 clusters were re-annotated. Trajectory inference was initially performed using

Slingshot[18] based on the recomputed UMAP embedding, and then utilized STREAM (v1.1)[19] to reconstruct gene expression matrix in a structure called the principal graph, which is a set of curves that naturally describe the cell-pseudotime, trajectories, and branching points. In detail, 'seed_elastic_principal_graph (clustering="kmeans", n_clusters=12, use_vis=True)' was initially applied to construct the principal graph based on recomputed UMAP embedding, and then 'elastic_principal_graph (epg_alpha=0.02, epg_mu=0.1, epg_lambda=0.01)' was applied to smooth the trajectory. Two trajectories were identified: AT1 lineage and the AT2 lineage. Pseudotime-correlated genes were visualized using 'SCP::GroupHeatmap()'. PAGA analysis[20, 21] was used to verify trajectory robustness, showing consistency with STREAM-derived trajectories.

### Supplementary Figure Legends

**Supplementary Fig. 1 | Preparation and characterization of stiffness-tunable GelMA hydrogels**

**a**, Schematic showing GelMA hydrogel synthesis with varying degrees of methacrylaiton (MA) (30%, 60%, 90%) to produce hydrogels of different stiffness (G30, G60, G90) through UV crosslinking. Long blue curve is Gelatin; Short red curve represents MA.

**b**, Frequency-dependent dynamic rheology measurements of storage modulus (G') and loss modulus (G") for G30, G60 and G90 hydrogels. Stiffness increases with MA modification degree.

**c**, Stress-strain curves showing mechanical behavior of the GelMA hydrogels. The inset magnifies the strain range of 0.050–0.100, highlighting the initial part where the stress and strain exhibit a linear increase.

**d**, Elastic modulus of G30, G60 an G90 hydrogels. Higher MA yields significantly stiffer hydrogels.

**e**, Swelling ratios of G30, G60 and G90 hydrogels. Insets show representative images of hydrogels post-swelling, indicating reduced swelling with increasing stiffness.

**f**, Scanning electron microscopy (SEM) images of hydrogels cross-section pore structures under varying stiffness. Lower stiffness (G30, i, iv) displays larger and loosely crosslinked pores, while higher stiffness (G90, iii, vi) shows smaller, denser pores.

**g**, Quantification of pore area for G30, G60 and G90 hydrogels , showing a significant reduction in pore size with increasing stiffness.

**h,** Degradation testing of hydrogels immerged in pH 7.4 PBS with 20 μg/mL collagenase type II. The hydrogel droplets were taken photos over time, showing size reduction due to degradation. Dashed circles indicate hydrogel boundaries.

**i**, Quantification of hydrogel hemisphere diameters during degradation. Higher stiffness (G90) leads to slower degradation compared to softer hydrogels (G30). Yellow, orange, and red denote G30, G60, and G90 stiffness groups, respectively. Statistical significance is indicated by * indicating comparison to G30, and # denoting comparisons to G60.

Data are presented as mean ± SEM, with individual dots representing values from 3–5 replicates in 3 independent experiments. $p$ values were calculated using one-way ANOVA and two-way ANOVA with Tukey's multiple comparison test. ## $p < 0.01$, *** and ### $p < 0.001$.

**Supplementary Fig. 2 | Hydrogel stiffness does not affect cell proliferation, cell cycle and apoptosis.**

**a**, Schematic illustrating the embedding AFE spheroids into GelMA hydrogels with different stiffness (G30, G60, G90), followed by UV crosslink to solidify the droplets and culture in LPC medium for 7 days.

**b**, Representative bright-field images showing LPC differentiated in varying stiffness hydrogels appeared epithelium structure with lumen inside .

**c**, Quantification of LPCs sizes across G30, G60, and G90 groups, showing no significant differences.

**d**, Representative live (green) and dead (red) staining of LPCs in G30, G60, and G90 hydrogels.

**e**, Quantification of the percentage of dead cells in different stiffness conditions, showing no significant variation.

**f**, Representative flow cytometry plots for propidium iodide (PI) staining, assessing cell viability in LPCs cultured under varying stiffness hydrogels.

**g**, qPCR analysis of *MKI67* mRNA expression, indicating no significant differences in cell proliferation among three groups.

**h**, Flow cytometry analysis of MKI67-positive LPCs showing similar proliferation rates across G30, G60, and G90 hydrogels.
**i**, qPCR analysis of proliferation-related genes (*AURKA* and *PCNA*) showing no stiffness-dependent changes.
**j**, qPCR analysis of cell cycle-related genes (*CDK4*, *CCNB1*, and *CCNE1*) showing comparable expression levels across stiffness groups.
**k,** qPCR analysis of apoptosis-related genes (*CASP3*, *MYBL2*, and *CDKN1A*) demonstrating no significant induction of apoptosis under varying stiffness.
Data are presented as mean ± SEM, with individual dots representing as the individual values of 3 to 5 replicates with 3 repeats in each experiment. Statistical comparisons were performed using one-way ANOVA with Tukey's multiple comparison test..

**Supplementary Fig. 3 | Flow cytometry analysis of airway epithelial cell markers under varying stiffness.**
**a**, CC10 (SCGB1A1, secretory cells) shows reduced expression with increasing stiffness.
**b**, MUC5AC (goblet cells) exhibits a stiffness-dependent increase, particularly in G90.
**c**, FOXJ1 (ciliated cells) shows modest upregulation under G60 and G90.
**d**, KRT5 (basal cells) is significantly upregulated in G90 hydrogel.

**Supplementary Fig. 4 | TEM images of microvilli and tubular myelin in mature AT2 cells.**
**a**, TEM images showing microvilli (MV, green arrows) on the surface of AT2 cells in hALOs under G30, G60, and G90, showing no stiffness-dependent changes.
**b**, TEM images displaying tubular myelin (TM, purple arrow), a key marker of AT2 surfactant secretion, was observed in G60 and G90 hALOs but not in G30.
**Supplementary Fig. 5 | single-cell RNA seqencing (scRNA-seq) reveals stiffness-dependent regulation of airway epithelial differentiation.**
**a**, UMAP visualization of scRNA-seq data, showing cell clustering by stiffness (left panel, G30, G60, G90) and annotated cell types (right panel, mesenchyme, epithelium and neurons).
**b**, Feature plots highlighting mesenchymal, epithelial, and neuronal marker expression across UMAP clusters.
**c**, Heatmap of marker genes expression across cell types demonstrating the specificity (left) and consistency (right) of genes expression profiles across stiffness.
**d, e**, UMAP visualization (**d**) and violin plots (**e**) showing increased goblet cells with increasing stiffness.
**f**, **g**, UMAP visualization (**f**) and violin plots (**g**) of secretory1 cells, which decreased with increasing stiffness.
**h**, **i**, UMAP visualization (**h**) and violin plots (**i**) showing Secretory2 cells downregulated under higher stiffness.
**j**, Venn diagram of downregulated genes across stiffness comparisons (G90-G60, G90-G30, G60-G30), identifying shared stiffness-responsive genes.
**k**, Heatmap of 353 differentially expressed genes (DEGs) down-regulated genes, including stress-response genes (*HSPA1A*, *HSP90AA1*, *HSBP1*) and secretory cell markers (*SCGB3A2*).
**l**, Pathway enrichment analysis of down-regulated genes, highlighting pathways such as cytoplasmic translation, oxidative phosphorylation, Wnt signaling regulation, and cellular stress response.
**m**, qPCR analysis of relative gene expression levels of *MUC5AC*, *SCGB1A1*, and *SCGB3A2* in hAWOs

treated with the combination of TRULI and FG-4592. ***$p < 0.001$, analyzed by one-way ANOVA with Dunnett's multiple comparison test (n=3).

**n**, Comparison of the expression levels of *SCGB3A2* under the treatments of TRULI and FG-4592. ***$p < 0.001$, analyzed by one-way ANOVA with Dunnett's multiple comparison test (n=3).

**o**, Gene expression of *MUC5AC*, *SCGB1A1*, and *SCGB3A2* under the individual treatments CHIR-99021. **$p < 0.01$, ***$p < 0.001$, analyzed by unpaired t test, one-way ANOVA with Dunnett's multiple comparison test (n=3).

**Supplementary Fig. 6 | scRNA-seq elucidates ECM stiffness-dependent alveolar differentiation in hALOs.**

**a,** UMAP visualization of scRNA-seq data from hALOs differentiated under varying stiffness GelMA (G30, G60, G90), showing clustering of epithelial, mesenchymal, and neuronal cell types.

**b**, Feature plots displaying the expression of epithelial, mesenchymal, and neuronal marker genes across pseudotime trajectories.

**c**, Validation of curated annotated cell types demonstrating the specificity (left) and consistency (right) of gens expression profiles across stiffness.

**d**, Heatmap depicting gene expression changes along the bifurcation lineages (Lineage 1 and Lineage 2) of hALOs differentiation. The left panel represented AT1lineage, including AT1-differentiation related genes such as *CLIC5*, *AGER* and *CAV1*, while the right panel represented AT2-lineage of genes such as *SFTPC*, *LAMP3*, *ABCA3*.

**e**, Partition-based Graph Abstraction (PAGA) trajectory analysis illustrating the differentiation relationships among AT1, proliferating AT1, proliferating AT2, AT2, and tip ETV5+ cells derived from hALOs. The thicker the line, the stronger the correlation of differentiation.

**f**, Venn diagram illustrating downregulated differential expressed genes (G90 vs G60, G90 vs G30, G60 vs G90) across stiffness variations, identifying shared and stiffness-specific genes. The key overlapped region shows 125 genes consistently downregulated with higher stiffness.

**g**, **h,** Heatmap displaying expression of shared down-regulated genes expression (**g**) accompanied by pathway enrichment analysis (**h**).

**Supplementary Fig. 7 | Schematic of ECM stiffness regulates epithelium cells fate during lung development.**

**a**, **LPC**: stiff hydrogel upregulates of LPCs markers (NKX2-1, SOX9, SOX2), with YAP and TGF-β signaling pathways notably influencing NKX2-1 expression.

**b**, **hAWOs**: high stiffness promotes key markers for proximal airway goblet, ciliated, and basal cells, while soft hydrogels favors the secretory cells emerged in the proximal-distal transition zone and distal airways. Notably, YAP, hypoxia, and Wnt signaling pathways regulate goblet and secretory cell expression.

**c**, **hALOs:** intermediate stiffness promotes alveolar maturation and the maturation of AT2 cell, while further increases stiffness triggers AT2 to AT1 transition, resulting in a higher proportion of AT1. Wnt and YAP signaling pathways are closely related to both alveolar maturation and alveolar epithelial cell proportions.

### Supplementary Figures

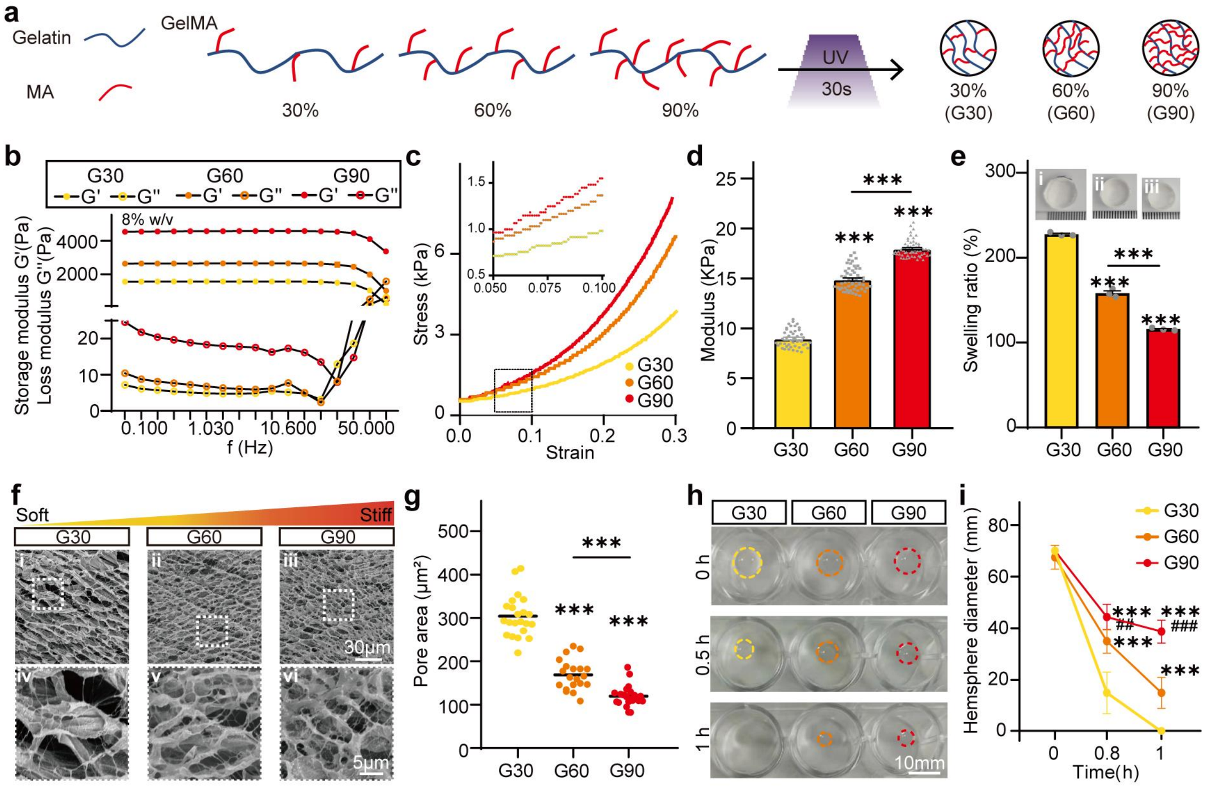


Supplementary Figure 1

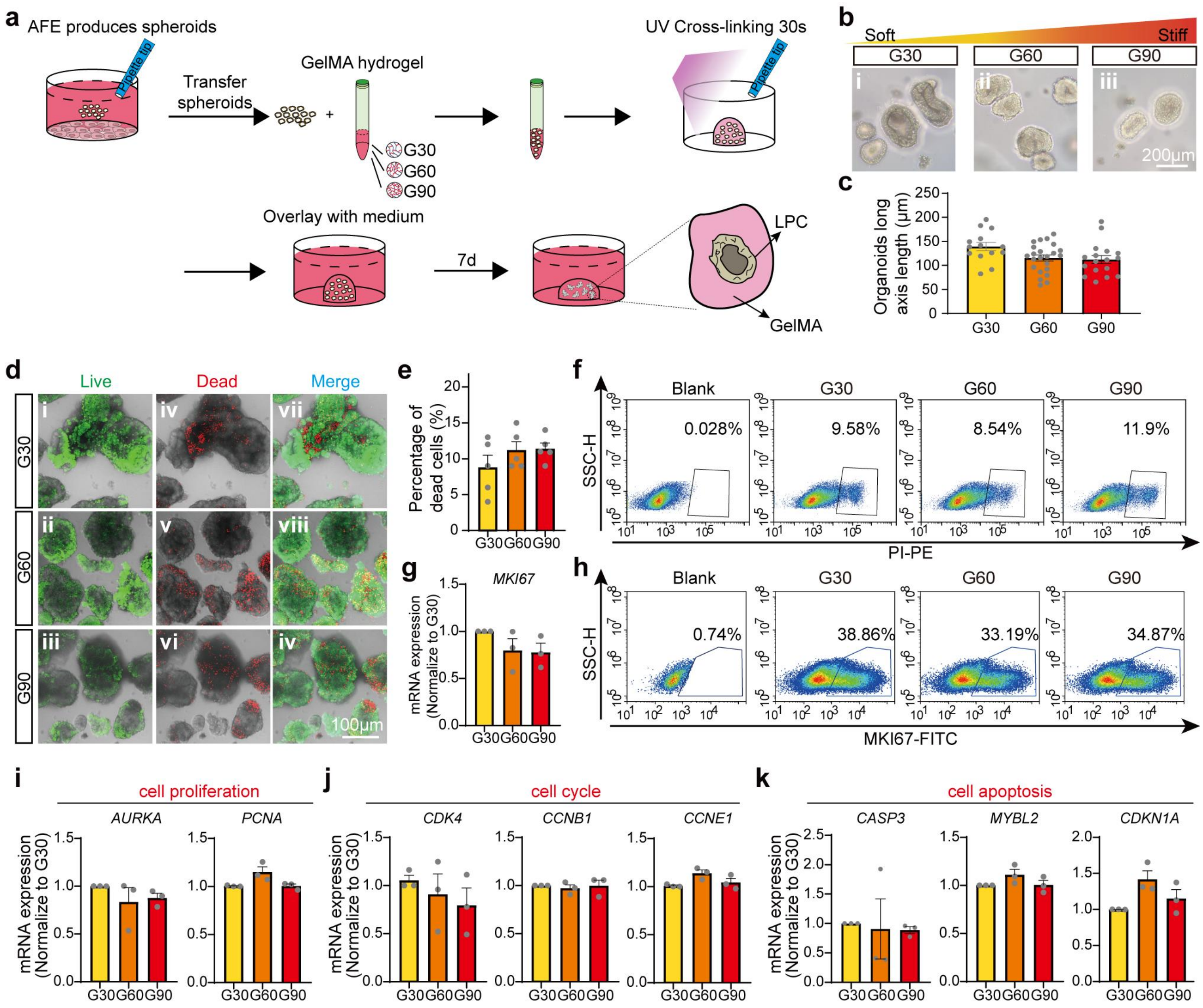




**Supplementary Figure 2**

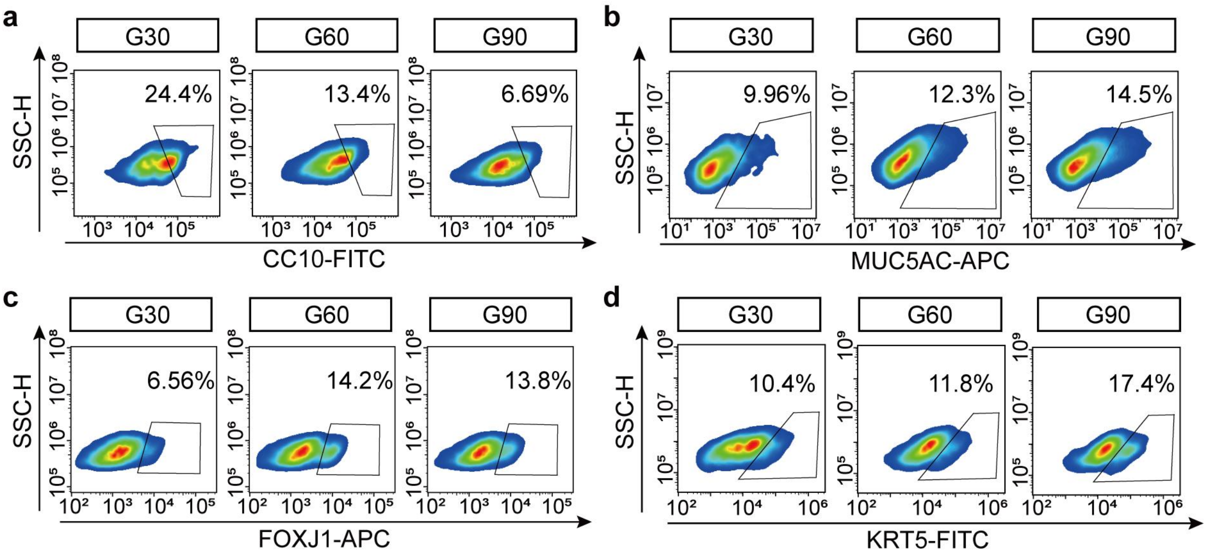


**Supplementary Figure 3**

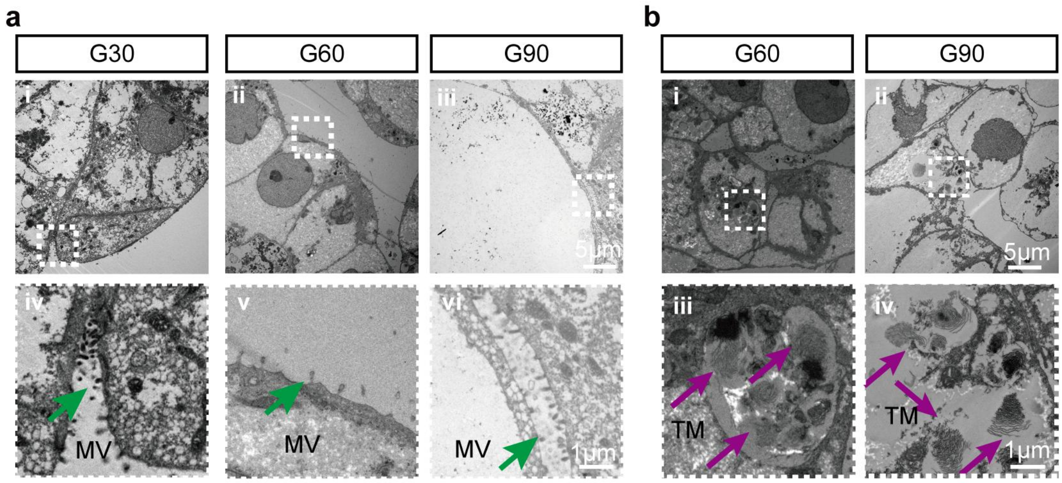


**Supplementary Figure 4**

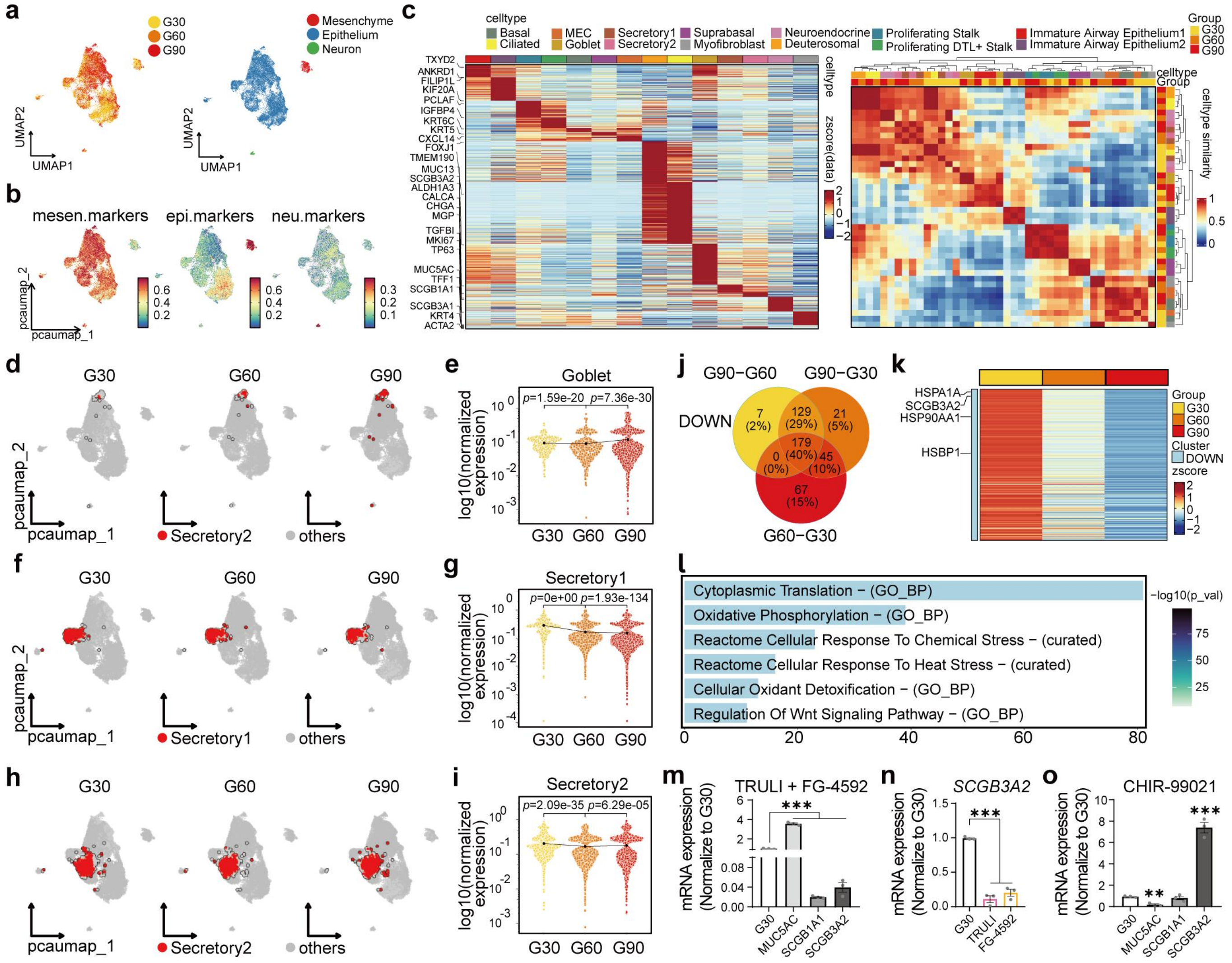


404

**Supplementary Figure 5**

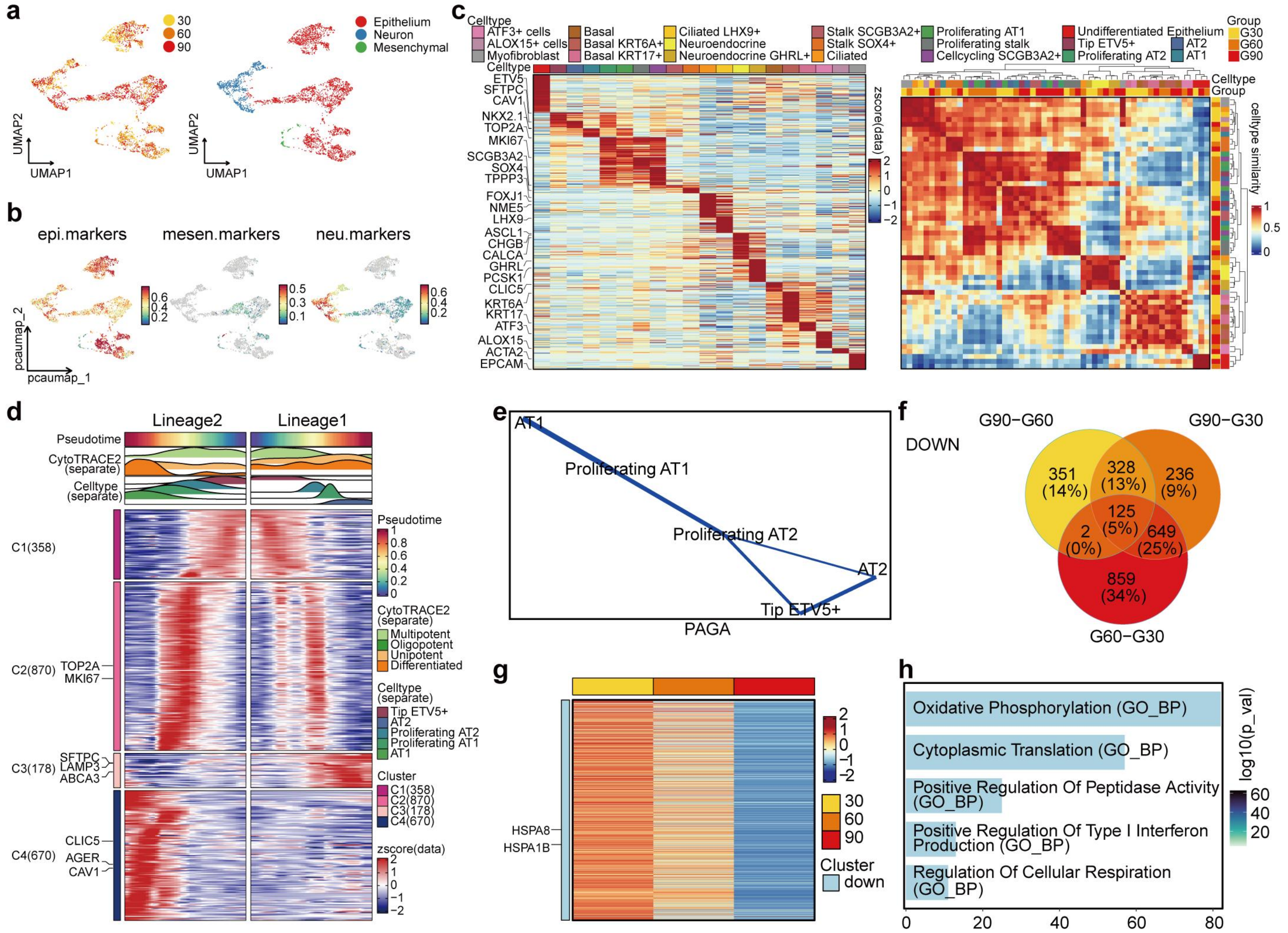


**Supplementary Figure 6**

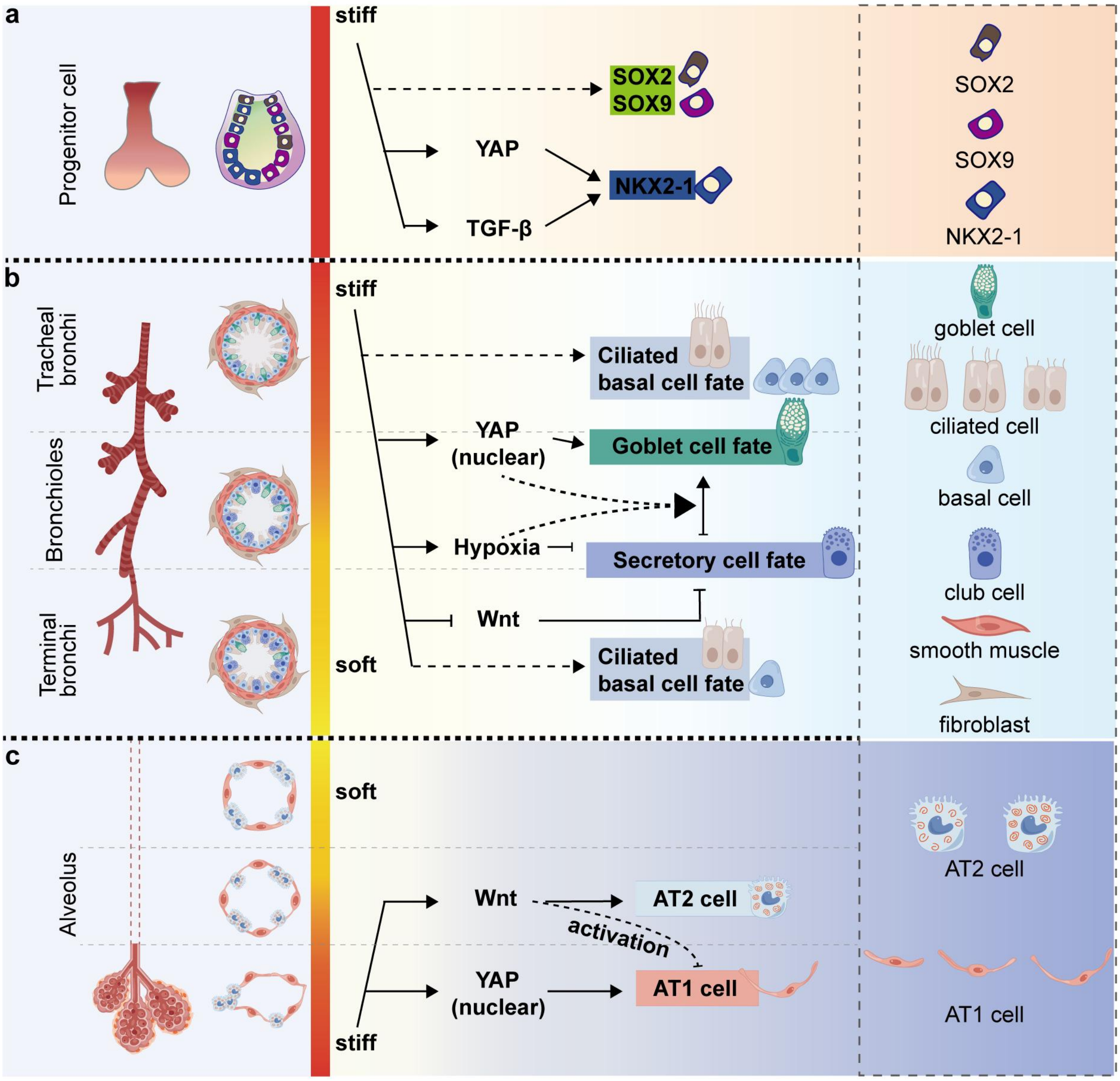


406 **Supplementary Figure 7**

**Supplementary Table 1. Primers used in the present study**

| Gene | Forward (5'-3') | Reverse (5'-3') |
|---|---|---|
| *NKX2-1* | CGGCATGAACATGAGCGGCAT | GCCGACAGGTACTTCTGTTGCTTG |
| *SOX2* | GCTTAGCCTCGTCGATGAAC | AACCCCAAGATGCACAACTC |
| *SOX9* | GACTACACCGACCACCAGAACTCC | CTGAGCTCGGCGTTGTG |
| *PAX8* | TGCCTCACAACTCCATCAGA | CAGGTCTACGATGCGCTG |
| *CDX2* | GGGCTCTCTGAGAGGCAGGT | GGTGACGGTGGGGTTTAGCA |
| *GATA4* | GCGGTGCTTCCAGCAACTCCA | GACATCGCACTGACTGAGAACG |
| *ID2* | GACAGCAAAGCACTGTGTGG | TCAGCACTTAAAAGATTCCGTG |
| *ABCA3* | CAGTGGACGGGCATACATCA | CCTTCAGCTGGGCGTAGAAA |
| *SFTPB* | TGCCTGGACCACCTCATCCTTG | GTCCTCACACTCTTGGCATAGG |
| *SFTPC* | AGCAAAGAGGTCCTGATGGA | CGATAAGAAGGCGTTTCAGG |
| *PDPN* | AGGAGAGCAACAACTCAACGGGAA | TTCTGCCAGGACCCAGAGC |
| *HOPX* | GCCTTTCCGAGGAGGAGAC | TCTGTGACGGATCTGCACTC |
| *AGER* | GCCACTGGTGCTGAAGTGTA | TGGTCTCCTTTCCATTCCTG |
| *SARS-CoV-2 NP* | CTGCAGATTTGGATGATTTCTCC | CCTTGTGTGGTCTGCATGAGTTTAG |
| *MUC5AC* | ACCAATGCTCTGTATCCTTCCC | GTTTGGGTGGAGTAAGCCACA |
| *SCGB1A1* | TTCAGCGTGTCATCGAAACCC | ACAGTGAGCTTTGGGCTATTTTT |
| *MKI67* | GAAAGAGTGGCAACCTGCCTTC | GCACCAAGTTTTACTACATCTGCC |
| *AURKA* | GCAACCAGTGTACCTCATCCTG | AAGTCTTCCAAAGCCCACTGCC |
| *PCNA* | ACACTAAGGGCCGAAGATAACG | ACAGCATCTCCAATATGGCTGA |
| *CDK4* | CTTTGGCAGCTGGTCACATGG | CTCAGATCAAGGGAGACCCTCAC |
| *CCNB1* | GACCTGTGTCAGGCTTTCTCTG | GGTATTTTGGTCTGACTGCTTGC |
| *CCNE1* | TGGATGTTGACTGCCTTGAA | TCCCCGTCTCCCTTATAACC |
| *CASP3* | GGCATGGAGAACACTGAAAAC | GCGAATCTGTTTCTTTGCATG |
| *MYBL2* | CACCAGAAACGAGCCTGCCTTA | CTCAGGTCACACCAAGCATCAG |
| *CDKN1A* | CGATGGAACTTCGACTTTGTCA | GCACAAGGGTACAAGACAGTG |
| *SCGB3A2* | GGCTAAGGAAGTGTGTAAATGAGC | CCATCCACCTCCGCTCTTTATC |
| *GAPDH* | TGCACCACCAACTGCTTAGC | GGCATGGACTGTGGTCATGAG |

**Supplementary Table 2. Antibodies used in the present study**

**Primary antibodies**

| **Antigen** | **Species** | **Cat.No.** | **Company** | **Dilution** |
|---|---|---|---|---|
| NKX2-1 | Rabbit | Ab76013 | Abcam | 1:250 (IF) (FC)<br>1:2000 (WB) |
| SOX9 | Mouse | Ab76997 | Abcam | 1:500 (IF) (FC) |
| SOX9 | Goat | AF3075 | R&D Systems | 1:50 (IF) (FC) |
| SOX2 | Goat | Ab239218 | Abcam | 1:50 (IF) (FC) |
| SOX2 | Mouse | Sc-365823 | Santa Cruz Biotechnology | 1:100 (IF) (FC) |
| UGRP1 | Goat | AF3545 | R&D Systems | 1:500 (IF) |
| Uteroglobin | Rabbit | Ab40873 | Abcam | 1:300 (IF) (FC) |
| Uteroglobin (SCGB1A1) | Mouse | Sc-365992 | Santa Cruz Biotechnology | 1:200 (IF) (FC)<br>1:1000 (WB) |
| MUC5AC | Mouse | Ab3649 | Abcam | 1:400 (IF) (FC) |
| MUC5AC | Mouse | EM1707-36 | HUABIO | 1:1000 (WB) |
| KRT5 | Mouse | Sc-32721 | Santa Cruz Biotechnology | 1:100 (IF) (FC) |
| AcTub | Mouse | T7451 | Sigma-Aldrich | 1:400 (IF) |
| FOXJ1 | Mouse | 14-9965-82 | eBioscience/invitrogen | 1:500 (IF) (FC) |
| SFTPB | Mouse | Sc-133143 | Santa Cruz Biotechnology | 1:200 (IF) (FC)<br>1:1000 (WB) |
| SFTPC | Rabbit | WRAB-9337 | Seven Hills Bioreagents | 1:200 (IF) |
| AGER | Goat | AF1145 | R&D Systems | 1:250 (IF)<br>1:200 (WB) |
| SARS-CoV-2 NP | Rabbit | 40143-T62 | SinoBiological | 1:200 (IF) |
| MKI67 | Mouse | 558616 | BD Transduction Laboratories | 1:200 (FC) |
| ZO-1 | Mouse | 33-9100 | Abcam | 1:300 (IF) |
| E-cadherin | Goat | 610181 | BD Transduction Laboratories | 1:500 (IF) |
| E-cadherin | Goat | AF748 | R&D Systems | 1:100 (IF) |
| HOPX | Mouse | Sc-398703 | Santa Cruz Biotechnology | 1:250 (FC) |
| YAP | Mouse | Sc-101199 | Santa Cruz Biotechnology | 1:250 (IF) |
| β-tubulin | Mouse | AF2839 | Beyotime | 1:5000 (WB) |
| GAPDH | Mouse | AF5009 | Beyotime | 1:5000 (WB) |

**Secondary antibodies**

| **Antigen** | **Cat.No.** | **Company** | **Dilution** |
|---|---|---|---|
| Alexa Fluor 488 Donkey anti-Rabbit | A-21206 | Invitrogen | 1:500 (IF) |
| Alexa Fluor 488 Donkey anti-Goat | A-11055 | Invitrogen | 1:500 (IF) |

| | | | |
|---|---|---|---|
| Alexa Fluor 488 Donkey anti-Mouse | A-21202 | Invitrogen | 1:500 (IF) |
| Alexa Fluor 546 Donkey anti-Mouse | A-10016 | Invitrogen | 1:500 (IF) |
| Alexa Fluor 546 Donkey anti-Rabbit | A-10040 | Invitrogen | 1:500 (IF) |
| Alexa Fluor 546 Donkey anti-Goat | A-11056 | Invitrogen | 1:500 (IF) |
| Alexa Fluor 647 Donkey anti-Mouse IgG | A-31571 | Invitrogen | 1:500 (IF) |
| Donkey anti-Rabbit IgG-647 | Ab150075 | Abcam | 1:500 (IF) |
| Donkey anti-Goat IgG-647 | Ab150135 | Abcam | 1:500 (IF) |
| HRP-conjugated Donkey Anti-Goat IgG (H+L) | SA00001-3 | proteintech | 1:10000 (WB) |
| HRP-conjugated Goat Anti-Mouse IgG(H+L) | SA00001-1 | proteintech | 1:10000 (WB) |
| HRP-conjugated Goat Anti-Rabbit IgG(H+L) | SA00001-2 | proteintech | 1:10000 (WB) |
